# Spatiotemporal coordination of actin regulators generates invasive protrusions in cell-cell fusion

**DOI:** 10.1101/2024.10.25.620345

**Authors:** Yue Lu, Tezin Walji, Benjamin Ravaux, Pratima Pandey, Changsong Yang, Bing Li, Delgermaa Luvsanjav, Kevin H. Lam, Ruihui Zhang, Zhou Luo, Chuanli Zhou, Christa W. Habela, Scott B. Snapper, Rong Li, David J. Goldhamer, David W. Schmidtke, Duojia Pan, Tatyana M. Svitkina, Elizabeth H. Chen

## Abstract

Invasive membrane protrusions play a central role in a variety of cellular processes. Unlike filopodia, invasive protrusions are mechanically stiff and propelled by branched actin polymerization. However, how branched actin filaments are organized to create finger-like invasive protrusions is unclear. Here, by examining the mammalian fusogenic synapse, where invasive protrusions are generated to promote cell membrane juxtaposition and fusion, we have uncovered the mechanism underlying invasive protrusion formation. We show that two nucleation promoting factors (NPFs) for the Arp2/3 complex, WAVE and N-WASP, exhibit different localization patterns in the protrusions. While WAVE is closely associated with the plasma membrane at the leading edge of the protrusive structures, N-WASP is enriched with WIP along the actin bundles in the shafts of the protrusions. During protrusion initiation and growth, the Arp2/3 complex nucleates branched actin filaments to generate low density actin clouds, in which the large GTPase dynamin organizes the new branched actin filaments into bundles, followed by actin-bundle stabilization by WIP, the latter functioning as a novel actin-bundling protein. Disrupting any of these components results in defective protrusions and failed myoblast fusion in cultured cells and mouse embryos. Taken together, our study has revealed the intricate spatiotemporal coordination between two NPFs and two actin-bundling proteins in building invasive protrusions at the mammalian fusogenic synapse, and has general implications in understanding invasive protrusion formation in cellular processes beyond cell-cell fusion.

## Introduction

Actin-based invasive membrane protrusions are essential for many cellular processes, including cancer metastasis, leukocyte transendothelial migration, immunological synapse maturation, uterine-vulval attachment, and cell-cell fusion^1–5^. In cancer metastasis, actin-rich structures called invadopodia are required for cancer cell extravasation by invading and degrading the basement membrane^6,7^. Leukocytes utilize similar actin-rich structures called podosomes to invade endothelial cells during their transendothelial migration^1^. At the immunological synapses, cytotoxic T lymphocytes also generate actin-rich protrusions to deform the target cell surface to facilitate toxic protease release into the target cell^5^. During *C. elegans* development, a specialized uterine cell, anchor cell, uses invadopodia to breach the basement membrane and initiate uterine-vulval attachment^3^.

Cell-cell fusion is required for the development and regeneration of multicellular organisms^8–10^. Major insights into the mechanisms of cell-cell fusion came from studies of skeletal muscle cell fusion, in which mononucleated myoblasts fuse to form multinucleated, contractile muscle fibers^4,11–13^. Work in *Drosophila* myoblast fusion led to the discovery of the asymmetric fusogenic synapse, where an muscle cell (a fusion-competent myoblast) generates an actin-enriched podosome-like structure (PLS) to project multiple invasive protrusions into its fusion partner (a founder cell or a myotube) to promote membrane juxtaposition and fusion^14^. The physiological relevance of invasive protrusions in *Drosophila* myoblast fusion has been highlighted by fusion mutants in which the invasiveness of the protrusions is compromised^14–17^. Besides myoblast fusion, induced fusion between cultured *Drosophila* cells of hemocyte origin is also mediated by invasive protrusions, which bring cell membranes into close proximity for fusogen engagement^18^. These studies suggest that both muscle and non-muscle cells use invasive protrusions as a general mechanism in facilitating cell membrane fusion.

In *Drosophila*, invasive protrusions at the asymmetric fusogenic synapse are propelled by the Arp2/3 complex-mediated branched actin polymerization^13^. Two nucleation-promoting factors (NPFs) for the Arp2/3 complex, WASP and WAVE (also known as Scar), have redundant functions in activating actin polymerization at the fusogenic synapse^14^. WASP and WASP-interacting protein WIP (also known as Solitary or Sltr) colocalize with the F-actin core of the PLS^14,15,19^, whereas the localization of WAVE remains unclear. Genetic analyses in *Drosophila* showed that the WASP/WIP complex, but not the pentameric WAVE complex, is required for the invasiveness of the protrusions^14,15^. How WASP and WAVE, two NPFs with similar biochemical functions in activating the Arp2/3 complex, exert distinct functions in enhancing the invasiveness of protrusions is unclear.

To generate narrow invasive protrusions (∼250 nm diameter in *Drosophila*)^14^, the branched actin filaments must be organized into mechanically stiff bundles. Our recent study revealed that the large GTPase dynamin functions as a multi-filament actin-bundling protein required for *Drosophila* myoblast fusion^17^. Dynamin is enriched at the fusogenic synapse and promotes the invasiveness of membrane protrusions by bundling actin filaments while assembling into a helical structure^17^. A single *Drosophila* (or mammalian) dynamin helix can recruit and bundle 12 (or 16) actin filaments to the outer rim of the helix, making dynamin a highly efficient actin-bundling protein^17^.

Interestingly, once dynamin is assembled into a full helical structure, the latter undergoes rapid disassembly induced by GTP hydrolysis, releasing free dynamin dimers back into the cytosol and leaving behind loosely aligned 12 (or 16) actin filaments^17^. Despite the biochemical analyses of dynamin-mediated actin bundling, how dynamin spatially and temporally bundles actin filaments during invasive protrusion formation and whether additional actin crosslinkers are involved in stabilizing the loosely aligned actin filaments remain unknown.

While most of the current understanding of actin polymerization and bundling in myoblast fusion originated from *Drosophila* studies, our recent work on zebrafish fast muscle cell fusion revealed conserved molecular composition and cellular architecture at a vertebrate fusogenic synapse^20^. However, it remains to be determined whether a similar mechanism is underlying mammalian muscle cell fusion, an indispensable step in skeletal muscle development and regeneration^12^. During development, the somite-derived muscle progenitors first express the paired domain transcription factors, Pax3 and Pax7, which activate the myogenic regulatory factor (MRF) family of bHLH transcription factors, including MyoD, Myf5, MRF4/Myf6 and Myogenin (MyoG), leading to the differentiation and fusion of the muscle cells^21–24^. During adult muscle regeneration, muscle stem cells (satellite cells) are activated in response to injury and proliferate, differentiate, and fuse to repair injured myofibers^25,26^. To date, several actin cytoskeletal regulators have been implicated in mammalian myoblast fusion during development, such as N-WASP and its activator Cdc42^27,28^, an activator for the WAVE complex (Rac1)^28^, and the bi-partite guanine nucleotide exchange factor for Rac1 (Dock180 [also known as Dock1] and Elmo)^27–30^, or in C2C12 cells, such as a subunit of the WAVE complex (Nap1)^31^. In addition, two muscle-specific transmembrane fusogenic proteins, myomaker (MymK) and myomixer (MymX)/myomerger/minion, are required for myoblast fusion *in vivo* and can induce cell-cell fusion when co-expressed in fibroblasts^32–35^. Despite their fusogenic activity, MymK and MymX require a functional actin cytoskeleton to induce cell-cell fusion^32,35^, highlighting the importance of actin cytoskeletal dynamics in the fusion process. Despite the identification of actin regulators in myoblast fusion and the observed actin enrichment and/or membrane protrusions prior to the fusion of cultured muscle cells^36–39^, the localization of the actin regulators relative to the fusion sites and how these actin regulators build the fusion machinery have yet to be revealed.

In this study, we identified the mammalian fusogenic synapse and demonstrated a critical function for the F-actin-propelled invasive membrane protrusions in mammalian myoblast fusion. We revealed an intricate spatiotemporal coordination between two Arp2/3 NPFs, N-WASP and WAVE, and two actin-bundling proteins, dynamin and WIP, in building the invasive protrusions at the fusogenic synapse.

## Results

### Actin polymerization is upregulated prior to myoblast fusion

To assess the potential role for the actin cytoskeleton in mammalian myoblast fusion, we examined actin polymerization in muscle cells prior to fusion. Phalloidin staining of mouse C2C12 myoblasts at different time points after switching from growth media (GM; 10% fetal bovine serum) to differentiation medium (DM; 2% horse serum) revealed a gradual elevation of F-actin in muscle cells over time (Extended Data Fig. 1a). The increased amount of F-actin correlated with the enhanced expression of an MRF, MyoG, and a muscle structural protein, muscle myosin heavy chain (MHC) (Extended Data Fig. 1a,b), suggesting that actin polymerization correlated with muscle cell differentiation. Furthermore, overexpressing MyoG in a non-myogenic cell line, the human osteosarcoma U2OS cells, significantly increased the cellular F-actin content without changing the total actin level or transforming them into muscle cells (no muscle MHC expression or cell-cell fusion) (Extended Data Fig. 1c-e), demonstrating that actin polymerization is activated by MyoG independent of muscle cell differentiation. To examine the potential link between actin polymerization and myoblast fusion during muscle cell differentiation, we seeded the C2C12 cells on circular micropatterns to restrict a small number of cells within a microscopic field and followed them during the entire differentiation process using live cell imaging (Extended Data Fig. 1f). After two days in DM, we added SiR-Actin, a cell permeable and specific probe for F-actin^40^, to label F-actin for 30 min before imaging. Strikingly, only myoblasts with high SiR-Actin labeling underwent fusion, consistent with a role for the actin cytoskeleton in the fusion process (Fig. 1a,b and Supplementary Video 1). Taken together, these data suggest that actin polymerization is upregulated prior to myoblast fusion during myogenic differentiation.

**Figure 1.**
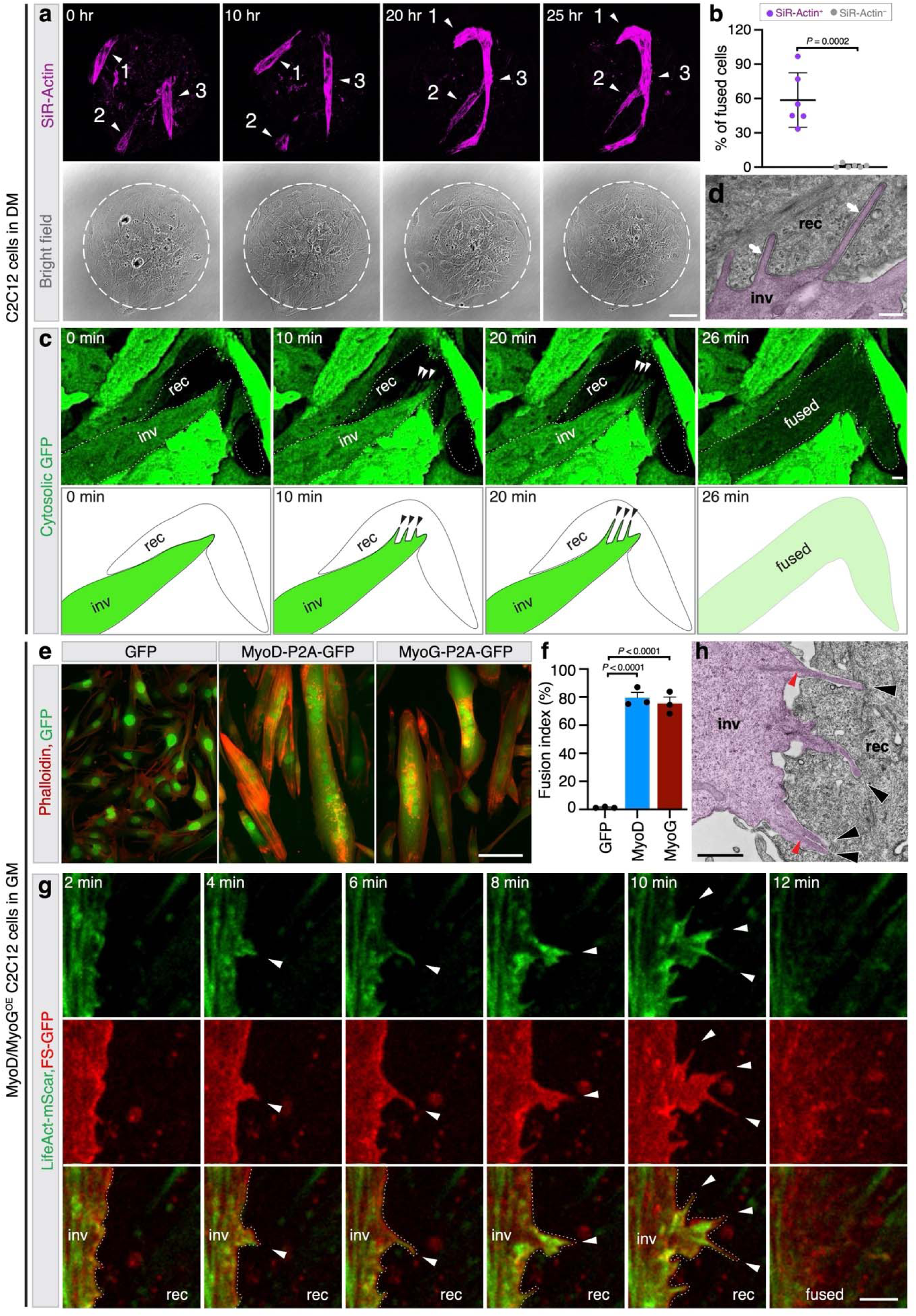
Mouse myoblast fusion is mediated by finger-like invasive protrusions at the fusogenic synapse. **a**, Still images of fusion events between SiR-Actin-labeled C2C12 cells cultured on a micropattern (schematic illustrated in **Extended Data** Fig. 1f). Three fusing myoblasts (arrowheads) are marked as 1, 2 and 3 (**Supplementary Video 1**). **b**, Quantification of the fusion of SiR-Actin^+^ vs. SiR-Actin^-^ cells tracked by live imaging. *n* = 6 independent experiments (**Statistics Source Data-Fig.1b)**. **c**, Still images of a fusion event between GFP^+^ and GFP^-^ C2C12 cells. Cell boundaries of the invading cell (inv) (arrowheads indicating the protrusions) and receiving cell (rec) are delineated by dotted lines except for the protrusive area (**Supplementary Video 2**). A schematic diagram of this fusion event is at the bottom. *n* = 22 fusion events were observed with similar results. **d**, An electron micrograph of C2C12 cells cultured for 48 hours in DM. The invading cell is pseudo-colored in magenta. Invasive protrusions at the cell-cell contact site are indicated by arrows. *n* = 15 cell-cell contact sites were observed with similar results. **e**, Phalloidin staining of C2C12 cells at low cell density in GM at 72 hours post-overexpression of GFP, MyoD-P2A-GFP, or MyoG-P2A-GFP. **f**, Quantification of fusion index for experiments in **e**. *n* = 3 independent experiments (**Statistics Source Data-Fig.1f)**. **g**, Still images of a fusion event between two MyoD^OE^ cells (schematic and zoomed-out view shown in **Extended Data** Fig. 1j and 1k, respectively). Cell boundaries are delineated by dotted lines in merged channels (**Supplementary Video 6**). *n* = 51 fusion events were observed with similar results. **h**, An electron micrograph of MyoD^OE^ cells at 48 hours post MyoD overexpression. The invading cell is pseudo-colored in magenta. Invasive protrusions and dense actin bundles along the shaft of the protrusions are indicated by black and red arrowheads, respectively. *n* = 13 cell-cell contact sites were observed with similar results. (**a**,**c**,**g**) Max z-projection of 8-10 focal planes from ventral plasma membrane (z-step size: 500 nm). (**e**) Epifluorescence images. (**g**) Single focal plane confocal images. Mean ± s.d. values are shown in the dot plot (**b**) and bar-dot plot (**f**), and significance was determined by two-tailed student’s t-test. Scale bars: 55 μm (**a**), 5 μm (**c**), 500 nm (**d**), 100 μm (**e**), 3 μm (**g**), and 2 μm (**h**).

### Myoblast fusion is mediated by invasive membrane protrusions

To observe the actin cytoskeletal dynamics at the sites of mammalian myoblast fusion, we performed live imaging of co-cultured C2C12 cells with or without cytosolic GFP at day two in DM. GFP^+^ finger-like membrane protrusions were observed projecting from invading cells into GFP^-^ fusion partners, the receiving cells, followed by cell fusion indicated by GFP transfer (Fig. 1c and Supplementary Video 2). In addition, live imaging of fusion between F-tractin-mCherry-expressing C2C12 cells with or without cytosolic GFP revealed F-actin enrichment within the invasive finger-like protrusions immediately before GFP transfer between the fusing cells (Extended Data Fig. 1g and Supplementary Video 3). Transmission electron microscopy (TEM) analyses of differentiating C2C12 cells confirmed that the protrusions at muscle cell contact sites were indeed invasive (Fig. 1d). Besides C2C12 cells, we performed live imaging of satellite cells in DM, as well as satellite cells induced to fuse by an Erk1/2 inhibitor (SCH772984) in GM^39^, and observed actin-propelled invasive protrusions at the fusion sites in both cases (Extended Data Fig. 1h,i and Supplementary Video 4,5).

Despite the observation of invasive protrusions at the fusion sites in C2C12 and satellite cells, the former need to reach confluency to fuse and latter are significantly smaller (∼1/3 of the size of C2C12 cells), making visualization of the fine actin structures at the fusion sites difficult in both cases. Strikingly, we found that overexpressing another MRF, MyoD, or MyoG in sparsely localized C2C12 cells in GM induced efficient myoblast fusion (Fig. 1e,f). Moreover, live imaging of MyoD-overexpressing (MyoD^OE^) C2C12 cells (will be referred to as MyoD^OE^ cells hereafter) revealed dynamic extension of multiple F-actin-enriched protrusions from the invading cells into the receiving cells (Fig. 1g, Extended Data Fig. 1j,k; 4-10 min and Supplementary Video 6) confirmed by TEM analysis (Fig. 1h). The F-actin in the invasive protrusions contained three actin isoforms, the sarcomeric α-actin and the cytosolic β- and γ-actin (Extended Data Fig. 2d). Only α-actin exhibited overall increased expression in differentiating C2C12 cells and MyoD^OE^ cells in GM (Extended Data Fig. 2a-c), but β- and γ-actin were highly enriched in the invasive protrusions. At the completion of fusion, the F-actin-enriched invasive structure rapidly dissolved (Fig. 1g and Extended Data Fig. 1k, 12 min). Taken together, our data demonstrate that F-actin-propelled invasive protrusions mediate mammalian myoblast fusion, and we will refer to the sites of mammalian myoblast fusion as the asymmetric fusogenic synapse hereafter. Since MyoD^OE^ cells in GM utilized similar invasive protrusions at the fusion sites as C2C12/satellite cells in DM and Erk1/2 inhibitor-treated satellite cells in GM, we used these cells for subsequent imaging analyses due to their large size and sparsely localized invasive protrusions.

### Branched actin polymerization promotes mammalian myoblast fusion

To ask what type of actin nucleators are involved in actin polymerization during mammalian myoblast fusion, we pharmacologically inhibited overall actin polymerization by cytochalasin D (CytD)^41^, the Arp2/3 complex by CK666^42^, or formins (nucleating linear actin filaments)^43^ by SMIFH2^44^, in differentiating C2C12 cells from day two to three, and observed the resulting myoblast fusion phenotype (Extended Data Fig. 3a,b). While all three drugs did not affect myoblast differentiation, myoblast fusion was significantly inhibited by CytD and CK666, but not SMIFH2 (Extended Data Fig. 3b-d), suggesting a role for branched, instead of linear, actin polymerization in mouse myoblast fusion. Since the Arp2/3 complex is activated by NPFs^45^, including WASP and WAVE, we tested their potential roles in mouse myoblast fusion. The mouse genome encodes multiple WASP and WAVE family members^46^, among which N-WASP (and one of the WASP-interacting proteins, WIP2) and WAVE2, in addition to Arp2/3, were expressed in both C2C12 cells and satellite cells at similar levels in GM and DM (Extended Data Fig. 3e,f). Due to their expression in muscle cells, we focused our subsequent analyses on N-WASP, WIP2, WAVE2, and the Arp2/3 complex.

Muscle differentiation involves two distinct but related processes – the expression of MRFs/muscle structural proteins, and the fusion between mononucleated myoblasts to generate multinucleated myofibers^12^. Inhibition of MRF expression could result in a defect and/or delay in myoblast fusion^47^. Since N-WASP, WIP2, WAVE2, and Arp2/3 were expressed in muscle cells before and during differentiation (Extended Data Fig. 3e,f), it is possible that they play a role in both muscle-specific gene expression and myoblast fusion. Indeed, when we knocked down these genes in C2C12 cells several days before switching to DM, we observed a ∼3 day delay of MyoG expression (Extended Data Fig. 3g,i-k), which could contribute in part to the severe myoblast fusion defect observed on day five in these cells (Extended Data Fig. 3h,l). To uncouple the functions of these actin regulators in MRF expression vs. myoblast fusion, we used the doxycycline (Dox)-inducible shRNA system to conditionally knock down these genes in C2C12 cells without causing a significant delay in MyoG expression (Fig. 2a,b and Extended Data Fig. 3m). Conditional knockdown (cKD) of N-WASP, WIP2, WAVE2, or Arp2 all resulted in severe fusion defects (Fig. 2c,d), suggesting that these actin regulators are required for myoblast fusion.

**Figure 2.**
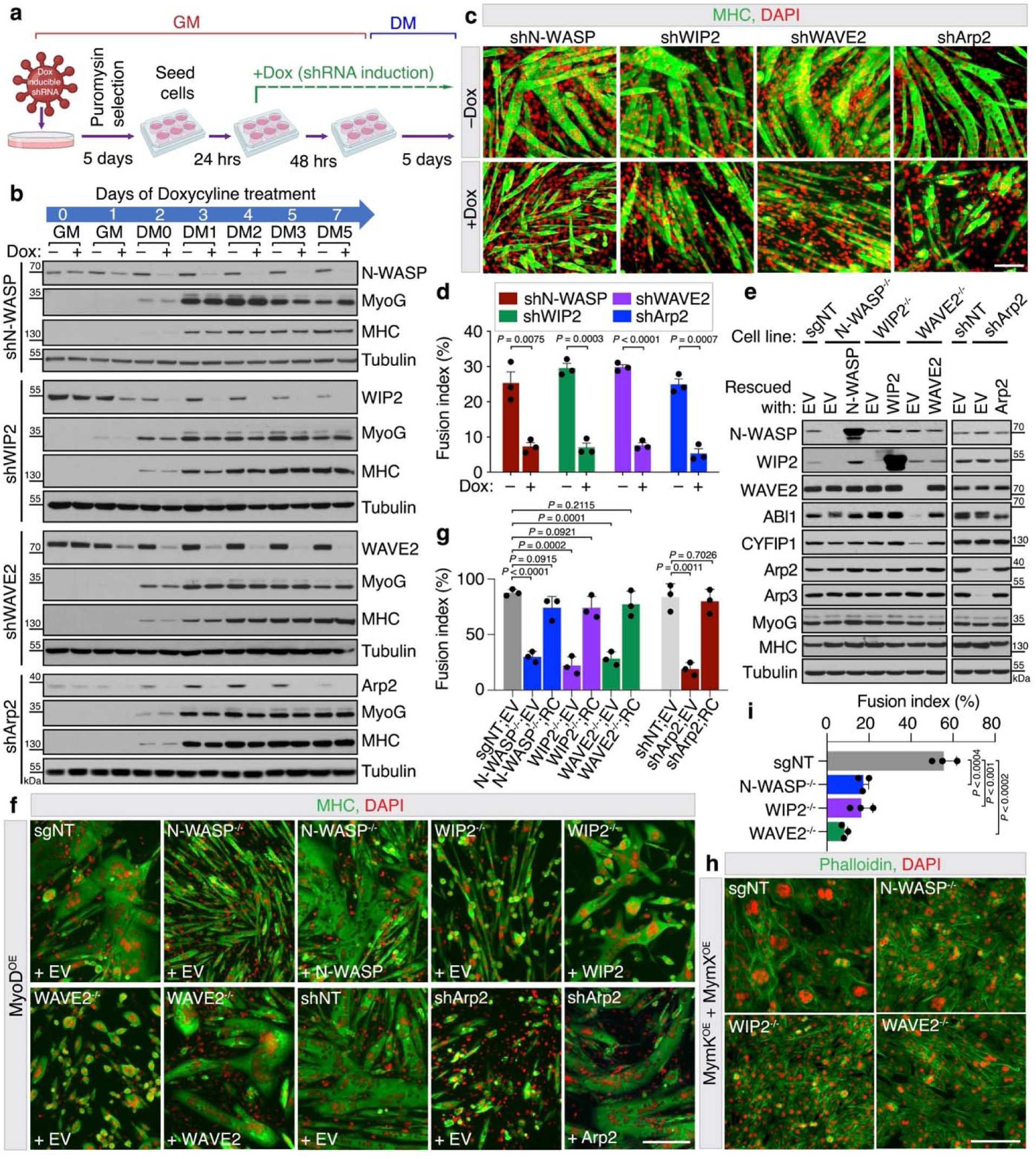
Branched actin polymerization promotes mouse myoblasts fusion. **a**, Schematic of conditional knockdown (cKD) of genes during myoblast fusion. **b**, Western blots showing the cKD of branched actin polymerization regulators during myoblast fusion. Control (-Dox) and KD (+Dox) cells as described in (**a**) were harvested at indicated time points to examine the protein levels of NPFs, the Arp2/3 complex, and muscle differentiation markers – MyoG and MHC. Quantification of MyoG and MHC protein levels at each indicated time point is shown in **Extended Data** Fig. 3m (**Statistics Source Data-Extended Data** Fig. 3m). *n* = 3 independent experiments. **c**-**d**, Immunostaining with anti-MHC in the control and cKD cells described in (**b**) at day five of differentiation. Quantification of fusion index is shown in (**d**) (**Statistics Source Data-Fig.2d**). **e**, Western blots showing that depletion of N-WASP, WAVE2, or Arp2, but not WIP2, led to a decrease in the expression of other members of the respective complexes. Note that in WIP2^-/-^ cells, N-WASP expression remained unchanged. In rescued cells, expression of other members of the protein complex recovered to normal level. Quantification of MyoG and MHC protein levels is shown in **Extended Data** Fig. 3n **(Statistics Source Data-Extended Data** Fig. 3n). *n* = 3 independent experiments. EV: empty vector, sgNT: non-targeting sgRNA, shNT: non-targeting shRNA. **f**-**g**, Immunostaining with anti-MHC in the cells described in (**e**) at 72 hours post MyoD overexpression. Quantification of the fusion indexes is shown in (**g**) (**Statistics Source Data-Fig.2g**). **h**-**i**, Phalloidin staining of control or NPF KO cells describe in (**e**) at two days post MymK and MymX co-overexpression in GM. Quantification of fusion index is shown in (**i)** (**Statistics Source Data-Fig.2i**). (**c**,**f**,**h**) Epifluorescence images. Mean ± s.d. values are shown in the bar-dot plot (**d**,**g**,**i**), and significance was determined by two-tailed student’s t-test. *n* = 3 independent experiments were performed. Scale bars: 100 μm (**c**) and 200 μm (**f**,**h**).

In addition to the cKD experiments, we generated N-WASP, WIP2 and WAVE2 knockout (KO) C2C12 cell lines. Deleting WAVE2 did not affect the expression of N-WASP/WIP2, and vice versa. Interestingly, while deleting WIP2 did not affect the expression of N-WASP, N-WASP KO abolished WIP2 expression (Fig. 2e), suggesting that N-WASP is stable in the absence of WIP2, but WIP2 requires N-WASP for its stabilization. Since all the KO cells did not survive well in DM, we took advantage of the MyoD^OE^ system to analyze their fusion in GM (Fig. 2f). The MyoD^OE^ KO cells also exhibited severe myoblast fusion defects (Fig. 2f,g), even though MyoG and MHC expression and cell mobility were not affected (Fig. 2e, Extended Data Fig. 3n,o and Supplementary Video 7). The fusion defects were rescued by expressing the corresponding knocked-out genes, demonstrating the specificity of the KOs (Fig. 2f,g). Since Arp2 KO cells were lethal, we generated Arp2 KD cells overexpressing MyoD in GM (Fig. 2e), which also exhibited a severe myoblast fusion defect (Fig. 2f,g).

Moreover, it has been reported that co-expressing the fusogenic proteins MymK and MymX in C2C12 cells in GM induced robust fusion^34^. Knocking out N-WASP, WIP2, or WAVE2 significantly inhibited MymK/MymX-induced C2C12 fusion in GM (Fig. 2h,i). It is worth noting that the levels of α-, β-, and γ-actin isoforms were unchanged in N-WASP, WIP2, and WAVE2 KD and KO cells, and only β-actin showed a moderate reduction in Arp2 KD cells (Extended Data Fig. 2e-h). Taken together, we have identified N-WASP, WIP2, WAVE2, and the Arp2/3 complex as the critical actin polymerization regulators required for mammalian myoblast fusion in addition to the fusogenic proteins.

### Distinct N-WASP/WIP2 and WAVE2 localization in the invasive structures

Given the requirement of N-WASP, WIP2, WAVE2, and the Arp2/3 complex in myoblast fusion, we asked whether these proteins are present at the fusogenic synapse. Live imaging of MyoD^OE^ cells revealed strong enrichment of mNeongreen (mNG)-tagged N-WASP, WIP2, and Arp2, but not WAVE2, with the F-actin-enriched structure at the fusogenic synapse prior to fusion, and their dissolution with F-actin when fusion was completed (Fig. 3a, Extended Data Fig. 4a-d and Supplementary Video 8). It was intriguing that WAVE2 was required for myoblast fusion, but mNG-WAVE2 was undetectable at the fusogenic synapse by live imaging. However, when we stained MyoD^OE^ cells with an anti-WAVE2 antibody, we observed WAVE2 enrichment in a thin layer at the leading edge of the invasive structure and at the protrusion tips, which was difficult to detect by live imaging (Fig. 3b,c and Extended Data Fig. 4e,f). In contrast, both N-WASP and WIP2 were more broadly localized and highly enriched along the shafts of the emerging protrusions, largely non-overlapping with WAVE2 (Fig. 3b,c, Extended Data Fig. 4f-h and Supplementary Video 9,10). Thus, even though both WAVE2 and N-WASP are known to be recruited to and activated at the plasma membrane^48–51^, they are enriched in different domains in the invasive structures (leading edge/tip vs. shaft), suggesting that they may play distinct roles in protrusion formation.

**Figure 3.**
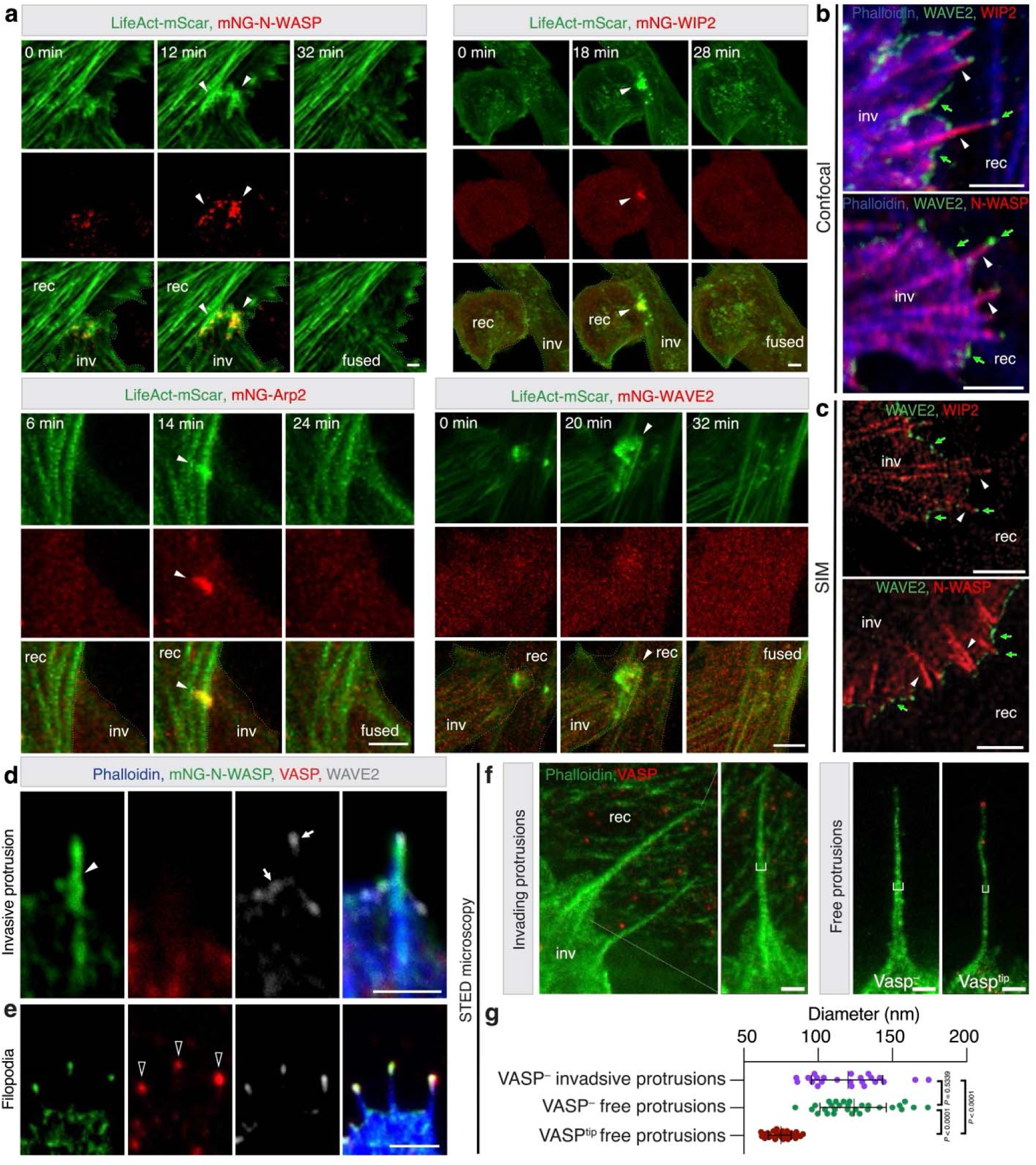
The localization of branched actin polymerization machinery in invasive protrusions. **a**, Still images of live imaging of stable C2C12 cell lines co-expressing LifeAct-mScar and mNeongreen (mNG)-N-WASP (or mNG-WIP2, mNG-WAVE2, or Arp2-mNG) after 48 hours of MyoD overexpression. Cell boundaries are delineated by dotted lines in the merged panels (**Supplementary Video 8**). *n* ≥ 15 fusogenic synapses (indicated by arrowheads) were imaged for each protein with similar results. **b**-**c**, Representative images of confocal microscopy (**b**) or structured illumination microscopy (SIM) (**c**) of C2C12 cell lines expressing 3×Flag-WIP2 or 3×Flag-N-WASP and immunostained with anti-WAVE2, anti-Flag, and phalloidin after 48 hours of MyoD overexpression. Single-channel images in (**b**) are shown in **Extended Data** Fig. 4f. WAVE2 enrichment are indicated by green arrows. N-WASP and WIP2 enrichment are indicated by arrowheads. *n* ≥ 20 cells were imaged for each panel with similar results. **d**-**e**, Immunostaining with anti-WAVE2, anti-VASP, and phalloidin in mNG-N-WASP-expressing C2C12 cells after 48 hours of MyoD overexpression. Note that the VASP^-^ protrusions had enriched N-WASP in the shafts (arrowheads), and enriched WAVE2 at the leading edge of the invasive structure and at the tips of protrusions (arrows) (**d**), whereas the VASP^tip^ filopodia (VASP enrichments indicated by hollow arrowheads) did not contain N-WASP in the shafts, but had N-WASP and WAVE2 enriched at the tips (**e**). *n* = 25 protrusions of each type were imaged with similar results. **f**-**g**, Representative images of stimulated emission depletion (STED) microscopy of C2C12 cells immunostained with anti-VASP and phalloidin at 48 hours post MyoD overexpression. Quantification of the protrusion diameters at the midpoint (indicated by brackets in (**f**) is shown in (**g**). Note that the VASP^-^ invasive protrusions and VASP^-^ free protrusions had similar diameters, which were significantly wider than that of the VASP^tip^ free protrusions. Mean ± s.d. values are shown in the dot plot, and significance was determined by two-tailed student’s t-test. *n* = 23, 29 and 30 (up to down) protrusions for each type of protrusions were quantified (**Statistics Source Data-Fig.3g**). (**a**) Max z-projection of 8-10 focal planes from the ventral plasma membrane (z-step size: 500 nm) (single focal plane images are shown in **Extended Data** Fig. 4d). (**b**-**f**) Single plane confocal images. Scale bars: 5 μm (**a**), 2 μm (**b**-**e**) and 500 nm (**f**).

### Invasive protrusions are distinct from filopodia

To date, the best-known actin-propelled finger-like cellular protrusions are filopodia, which are elongated by linear actin nucleators and elongation factors^52^. For example, the linear actin elongation factors, the Ena/VASP proteins (Ena/VASP, Mena and EVL), have been shown to be enriched at the tips of filopodia to promote their extension^52–57^. Thus, we asked whether invasive protrusions also employ Ena/VASP proteins for their elongation by staining the MyoD^OE^ cells with antibodies against these proteins. VASP was enriched at the tips of ∼44% protrusions (will be referred to as VASP^tip^ protrusions hereafter) in MyoD^OE^ cells (Extended Data Fig. 5a-d), among which ∼21% tips also contained EVL, whereas Mena was rarely observed at any protrusion tips (Extended Data Fig. 5e-g). In all VASP^tip^ protrusions, N-WASP/WIP2 and WAVEs were also enriched at the tips (Fig. 3e and Extended Data Fig. 5c,e,f). However, in protrusions without VASP at the tips (VASP^-^ protrusions), N-WASP/WIP2 appeared highly enriched along the shafts, whereas WAVE2 remained at the tips (Fig. 3d and Extended Data Fig. 5b,d). Notably, none of the Ena/VASP proteins were enriched at the tips of invasive protrusions at the muscle cell contact sites (Extended Data Fig. 5h), indicating that all invasive protrusions were VASP^-^ at the tips and N-WASP/WIP2^+^ along the shafts.

Consistent with this, previous work in *Drosophila* has shown that the WASP/WIP complex is required for the invasiveness of protrusions at the fusogenic synapse^14^. Morphologically, the VASP^-^ protrusions were thicker (122.5 ± 23.0 nm diameter) than VASP^tip^ protrusions (74.9 ± 7.8 nm diameter) visualized by stimulated emission depletion (STED) microscopy (Fig. 3f,g). Taken together, based on their different morphologies and molecular compositions, finger-like cellular protrusions fall into two categories – invasive protrusions and filopodia. Invasive protrusions are thicker (∼123 nm diameter), enriched with N-WASP/WIP2 along the shafts, and devoid of VASP at the tips, whereas filopodia are thinner (∼75 nm), enriched with VASP at the tips, and devoid of N-WASP/WIP2 along the shafts.

### WAVE2 and N-WASP/WIP2 exhibit different functions in invasive protrusion formation

To investigate the roles for WAVE2 and N-WASP/WIP2 in invasive protrusion formation, we performed live imaging of wild-type and NPF KO MyoD^OE^ cells. Wild-type cells exhibited dynamic actin cytoskeletal activities at the cell cortex, manifested by the formation of both lamellipodia and finger-like protrusions emanating from the leading edge of the lamellipodia (1.05 ± 0.19 protrusions/minute per 10 μm of cell cortex) (Fig. 4a1,b and Supplementary Video 11). Immunostaining showed that 51.8 ± 4.7% of these protrusions were VASP^-^ with N-WASP/WIP2 enrichment along the shafts (Fig. 4c,d,h). WAVE2^-/-^ cells barely generated any lamellipodia, coinciding with a significant decrease in the number of finger-like protrusions (0.23 ± 0.08 protrusions/minute per 10 μm of cell cortex) (Fig. 4a2,b and Supplementary Video 11), consistent with the previous finding that WAVE-mediated dendritic network in lamellipodia provides the source of actin filaments for finger-like protrusions^58^. Among the small number of protrusions generated in WAVE2^-/-^ cells, 49.6 ± 4.4% were VASP^-^ with N-WASP enrichment along the shafts (Fig. 4e,h). In contrast, N-WASP^-/-^ cells, in which both N-WASP and WIP2 were absent (Fig. 2e), formed lamellipodia (with WAVE2 at the leading edge) and generated significantly lower number of VASP^-^ protrusions (3.69 ± 0.4%, Fig. 4f,h), despite similar numbers of total finger-like protrusions as in wild-type cells (1.24 ± 0.1 protrusions/minute per 10 μm of cell cortex) (Fig. 4a3,b and Supplementary Video 11), suggesting that N-WASP and/or WIP2 are required for generating VASP^-^ invasive protrusions. Interestingly, WIP2^-/-^ cells, in which N-WASP was still present (Fig. 2e), did not form lamellipodia but generated significantly fewer VASP^-^ protrusions (3.4 ± 0.6%, Fig. 4h), and more VASP^tip^ protrusions (1 ± 0.03/minute per 10 μm of cell cortex) with WAVE2 and even N-WASP enriched at the tips (Fig. 4a4,b,g and Supplementary Video 11). The WIP2^-/-^ protrusions (mostly VASP^tip^) appeared more bendy than those in control cells indicated by the tortuosity index (Fig. 5i). Thus, WIP2 is required for N-WASP localization to the shafts and for enhancing the mechanical stiffness of the protrusions. Arp2 KD cells, in which both WAVE2- and N-WASP-mediated branched actin polymerization was significantly reduced, exhibited no lamellipodia or finger-like protrusions, despite the normal overall cytosolic actin stress fibers (Fig. 4a5,b and Supplementary Video 11). These results confirmed that branched actin polymerization is required for generating invasive protrusions and that WAVE2 and N-WASP/WIP2 have distinct functions in protrusion formation.

**Figure 4.**
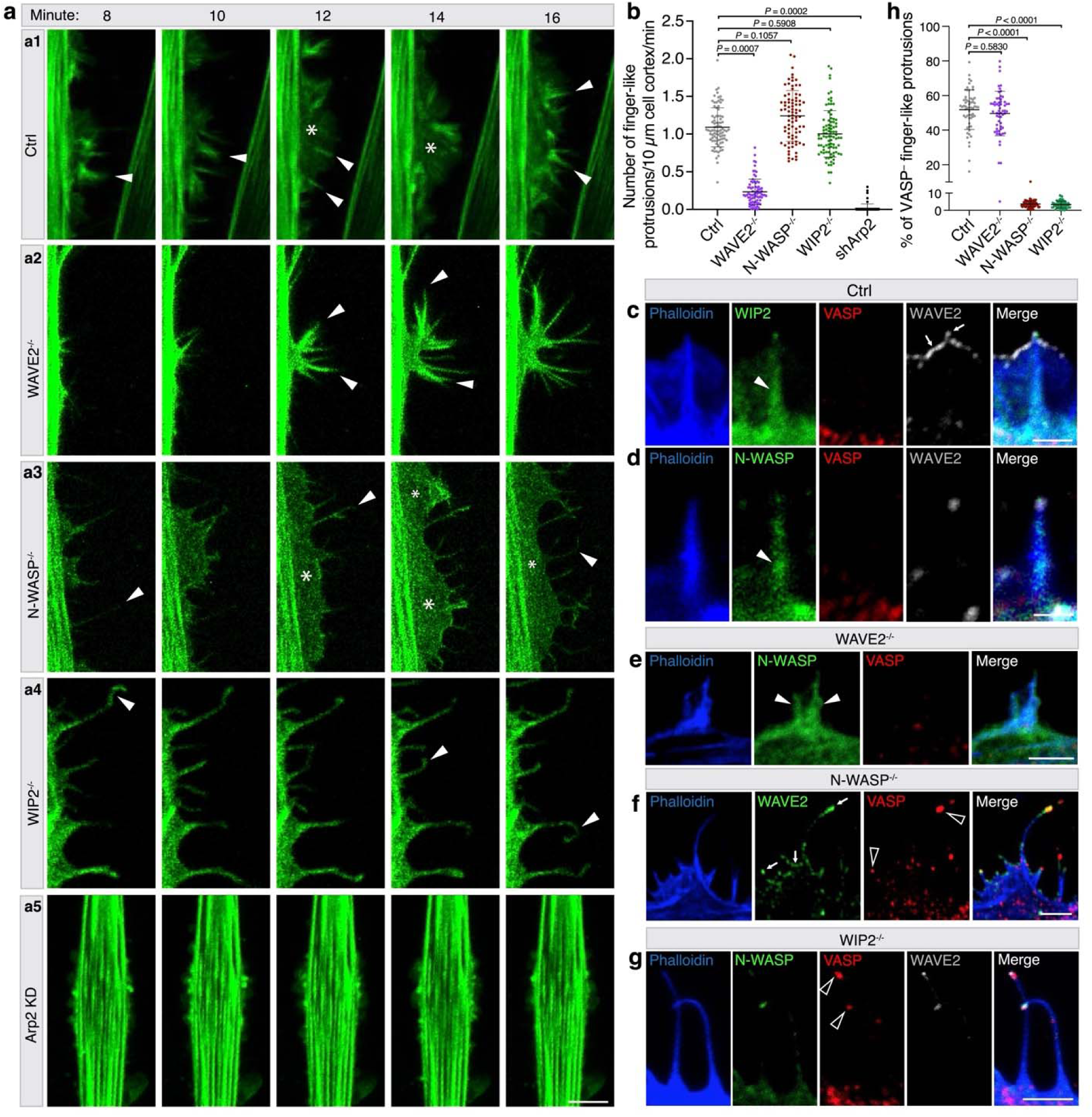
N-WASP/WIP2 and WAVE2 have distinct functions during protrusion formation. **a**, Still images of cortical protrusions of wild-type (Ctrl) (**a1**), NPF KOs (**a2**-**a4**), and Arp2 KD (**a5**) C2C12 cells at 48 hours post MyoD overexpression (**Supplementary Video 11**). Randomly selected cortical regions containing finger-like protrusions (arrowheads) and lamellipodia (asterisks) are shown. Note the straight vs. bendy finger-like protrusions projected from the leading edge of the lamellipodia in Ctrl vs. N-WASP^-/-^ cells; the loss of lamellipodia and the decreased number of protrusions in WAVE2^-/-^ cell; the bendy protrusions in WIP2^-/-^ cell; and no protrusions in Arp2 KD cells. **b**, Quantification of the number of newly formed finger-like protrusions in cells with the genotypes shown in (**a**). *n* = 86, 82, 90, 79 and 67 (left to right) cells from three independent experiments were measured for each genotype (**Statistics Source Data-Fig.4b**). **c**-**g**, Localization of N-WASP, WIP2, WAVE2, and VASP in individual protrusions of Ctrl and mutant MyoD^OE^ cells. The mNG-N-WIP2-expressing Ctrl (**c**), mNG-N-WASP-expressing Ctrl (**d**), mNG-N-WASP-expressing WAVE2^-/-^ (**e**), N-WASP^-/-^ (**f**), and mNG-N-WASP-expressing WIP2^-/-^ (**g**) cells were immunostained with anti-WAVE2, anti-VASP, and phalloidin at 48 hours post MyoD overexpression. **h**, Quantification of the percentage of VASP^-^ protrusions in cells with the genotypes shown in (**c**-**g**). The Ctrl, WAVE2^-/-^, N-WASP^-/-^, and WIP2^-/-^ cells were immunostained with anti-VASP and phalloidin at 48 hours post MyoD overexpression. *n* = 60 cells from three independent experiments were quantified for each genotype (**Statistics Source Data-Fig.4h**). (**a**) Max z-projection of 8-10 focal planes from the ventral plasma membrane (z-step size: 500 nm). (**c**,**g**) Single plane confocal images. Mean ± s.d. values are shown in the dot plot (**b**,**h**), and significance was determined by two-tailed student’s t-test. Scale bars: 10 μm (**a**), 3 μm (**e**,**f**,**g**), 2 μm (**c**) and 1.5 μm (**d**).

**Figure 5.**
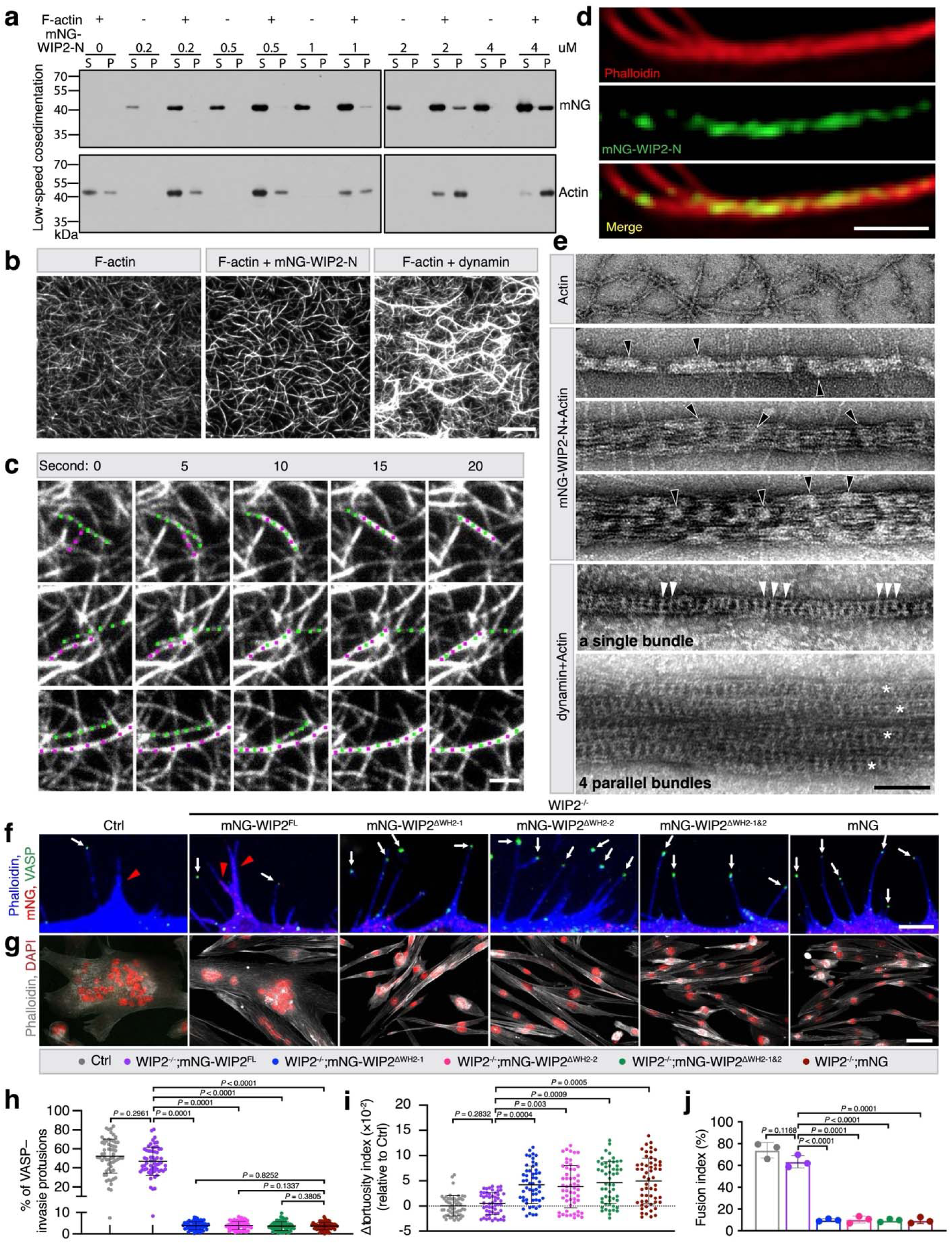
WIP2 is an actin-bundling protein. **a**, Low-speed co-sedimentation assay for F-actin and mNG-WIP2-N. **b**, TIRF images of WIP2- and Shi (*Drosophila* dynamin)-mediated actin bundling (**Supplementary Video 12**). **c**, Still images of time-lapse TIRF imaging showing three examples of mNG-WIP2-mediated actin filament bundling (**Supplementary Video 13**). The merging actin filaments are indicated by magenta or green dots. *n* = 3 independent imaging experiments were performed with similar results. **d**, SIM images of actin bundles assembled by Alexa Fluor 568-phalloidin (red) and mNG-WIP2-N (green). **e**, Electron micrographs of negatively stained actin filaments alone (top panel), WIP2-actin bundles (middle panels), and dynamin-actin bundles (bottom two panels). Three WIP2-actin bundles with different diameters are shown. A dynamin-actin single bundle and a dynamin-actin super bundle consisting of four parallel single bundles (each marked by an asterisk) are shown. A few mNG-WIP2-N patches are indicated by hollow arrowheads, and a few dynamin helical rungs are indicated by white arrowheads. **f**, Immunostaining with anti-VASP and phalloidin in wild-type cells (Ctrl), and WIP2^-/-^ cells expressing mNG-full-length (FL) WIP2, mNG-WIP2^ΔWH2-1^, mNG-WIP2^ΔWH2-2^, mNG-WIP2^ΔWH2-1&2^, or mNG after 48 hours of MyoD overexpression. The VASP^-^ and VASP^tip^ protrusions are indicated by arrowheads and arrows, respectively. Zoomed-out views of the cells are shown in **Extended Data** Fig. 6c. **g**, Phalloidin staining in cells with the genotypes shown in (**f**) at 72 hours post MyoD overexpression. **h**, Quantification of the percentage of VASP^-^ protrusions for each genotype shown in (**f**). *n* = 60 cells for each genotype from three independent experiments were analyzed. **i**, Quantification of the bendiness of the protrusions in each genotype shown in (**f**). *n* = 55 protrusions from three independent experiments for each genotype were analyzed. **j**, Quantification of fusion index of cells in (**g**). *n* = 3 independent experiments were performed. (**f**) Single plane confocal images. (**g**) Epifluorescence images. In (**a**,**b**,**c**,**d**,**e**), *n* = 3 independent experiments were performed with similar results. Mean ± s.d. values are shown in the dot plot (**h**,**i**) and bar-dot plot (**j**), and significance was determined by two-tailed student’s t-test (**Statistics Source Data-Fig.5h-j**). Scale bars: 10 μm (**b**), 2 μm (**c**), 1 μm (**d**), 100 nm (**e**), 5 μm (**f**) and 50 μm (**g**).

### WIP2 bundles F-actin and increases the mechanical strength of invasive protrusions

The strong enrichment of WIP2 with F-actin in the protrusion shafts (Fig. 4c and Extended Data Fig. 5b) is not surprising due to the presence of two actin-binding WH2 domains (WH2-1 and WH2-2) at the N-terminus of WIP proteins^19,59^. Using purified WH2-1, WH2-2, and WIP2-N (an N-terminal fragment containing both WH2-1 and WH2-2), we confirmed that both WH2 domains bound F-actin by high-speed co-sedimentation assays (Extended Data Fig. 6a). But to our surprise, low-speed co-sedimentation assays showed that WIP2-N bundled F-actin in a dosage-dependent manner (Fig. 5a).

In contrast, WH2-1 or WH2-2 alone did not show F-actin bundling activity (Extended Data Fig. 6b). Since both WH2 domains are required for WIP2’s actin-bundling activity, WIP2 appeared to function as a two-filament actin crosslinker. Using total internal reflection fluorescence (TIRF) microscopy, we observed that when WIP2-N was added to a dynamically polymerizing branched actin network, it rapidly “froze” the network by crosslinking actin and forming actin bundles at numerous locations, as did dynamin (Fig. 5b and Supplementary Video 12). At the single-filament level, WIP2 could induce bundling of two actin filaments, either connected or unconnected (Fig. 5c and Supplementary Video 13). SIM analysis revealed that WIP2 formed sparsely localized patches along the actin bundles (Fig. 5d). Consistent with this, negative-stain EM also showed light colored patches (formed by WIP2) along tight actin bundles (Fig. 5e).

Unlike the dynamin-mediated actin bundle, each of which is organized by a dynamin helix and has an average diameter of 32 nm^17^ (Fig. 5e), the WIP2-mediated actin bundles did not have a fixed diameter and exhibited various widths (Fig. 5e). These data demonstrate a function of WIP2 as an actin-bundling protein and suggest that it may do so in the shafts of invasive protrusions to make them mechanically stiff.

To investigate the requirement for the actin-bundling activity of WIP2 in invasive protrusions, we made internal deletions of one or both WH2 domains in the context of the full-length (FL) WIP2 protein, mNG-WIP2^△WH2-1^, mNG-WIP2^△WH2-2^, or mNG-WIP2^△WH2-1&2^, and tested their effects on VASP^-^ invasive protrusion formation in WIP2^-/-^ cells. While expressing mNG-WIP2^FL^ rescued the number of VASP^-^ protrusions from 3.6 ± 0.3% to wild-type level (46.9 ± 4.8%), deleting one or both WH2 domains failed to do so (Fig. 5f,h and Extended Data Fig. 6c). In addition, expressing mNG-WIP2^FL^, but not the mutant WIP2 proteins, rescued the bendiness of protrusions (from 4.9 ± 0.7 to 0.5 ± 0.3) (Fig. 5i) and fusion (from 9.6 ± 2.8% to 63.5 ± 5.7%) to wild-type levels (Fig. 5g,j). Thus, the actin-bundling activity of WIP2 mediated by both WH2 domains is required for invasive protrusion formation and myoblast fusion.

### Arp2/3-nucleated branched actin filaments are organized into bundles

The crucial role for the Arp2/3 complex and its NPFs in invasive protrusion formation prompted us to ask how newly polymerized branched actin filaments are organized into bundles. To observe the dynamics of actin polymerization during protrusion growth, we performed live imaging of MyoD^OE^ cells expressing LifeAct-mScar and Arp2-mNG (Supplementary Video 14). As shown in Fig. 6a, the Arp2/3 complex first induced the expansion of a lamellipodium-like structure on the side of a protrusion (Fig. 6a, 3-4 min). Since the density (unit fluorescence intensity) within the actin structure on the side was lower than that within the main protrusion, we will refer to the former as an “actin cloud” hereafter. Shortly after, a new F-actin bundle emerged from the actin cloud, coinciding with a decrease of Arp2 in the bundle (Fig. 6a, 6 min). Subsequently, new actin clouds continued to grow on the periphery of the new bundle (Fig. 6a, 7 min), eventually leading to bundle growth/thickening (Fig. 6a, 8 min).

**Figure 6.**
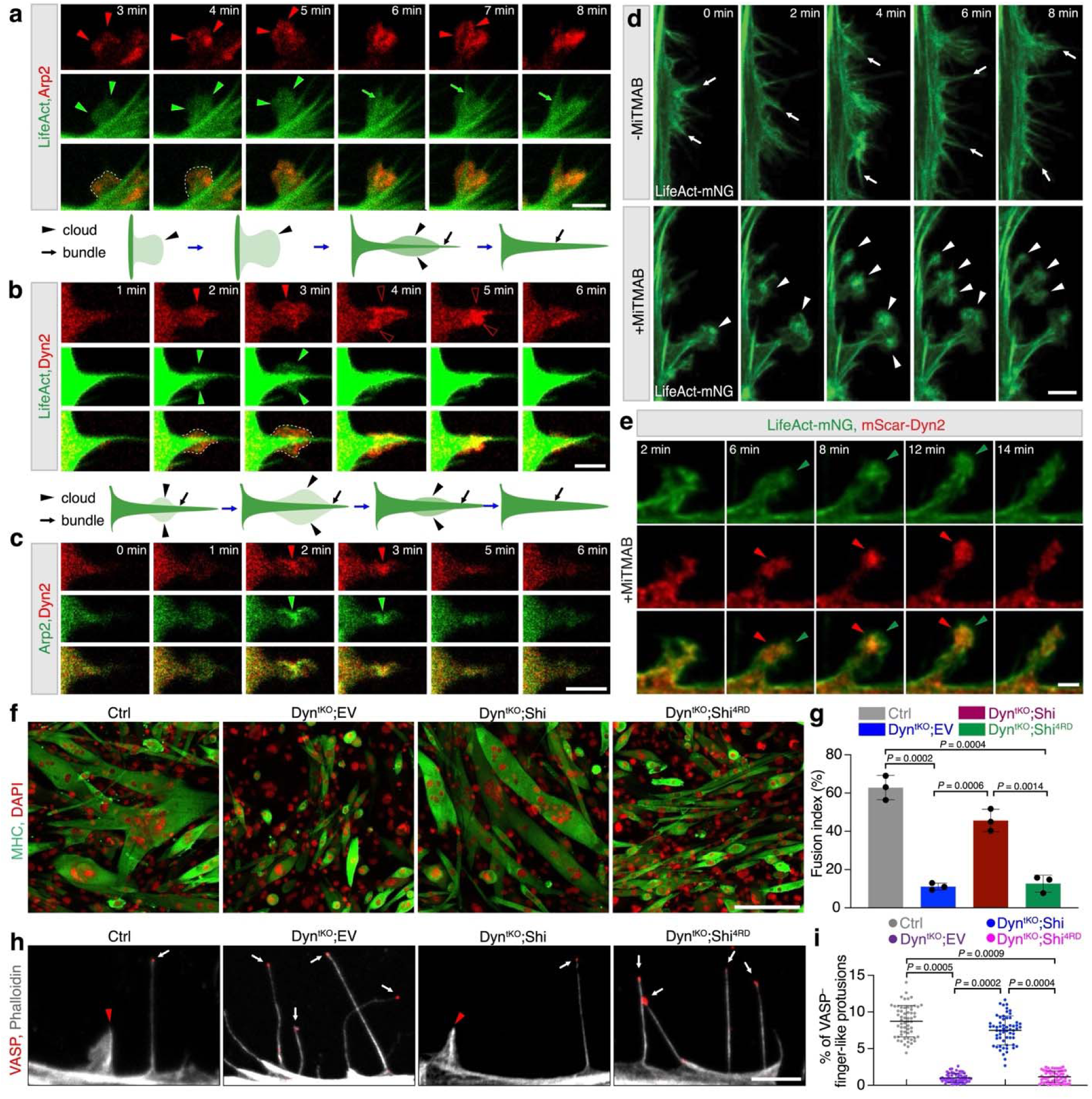
Dynamin is required for bundling the Arp2/3 complex-nucleated branched actin filaments. **a-c**, Dynamic enrichment of Arp2 and Dyn2 with newly polymerized branch actin filaments in C2C12 cells at 48 hours post MyoD overexpression (**Supplementary Video 14-16**). A schematic diagram is shown at the bottom of each panel of still images. *n* = 3 independent experiments were performed with similar results. Actin “clouds” are outlined by dotted lines in the merged channel in (**a**,**b**). In (**a**), Arp2-mScar enrichment, actin clouds, actin bundles are indicated by red arrowheads, green arrowheads and green arrows, respectively. In (**b**), at 2 and 3 min, actin clouds and mScar-Dyn2 enrichment are indicated by green and red arrowheads, respectively. At 4 and 5 min, mScar-Dyn2 enrichment is indicated by hollow arrowheads. In (**c**), mScar-Dyn2 Arp2-mNG enrichment are indicated by red and green arrowheads, respectively. **d**, Still images of cortical protrusions of LifeAct-mNG-expressing C2C12 cells treated without (top) or with (bottom) 5 μM MiTMAB, at 48 hours post MyoD overexpression (**Supplementary Video 17**). *n* = 3 independent experiments were performed with similar results. **e**, Localization of Dyn2 in the persistent actin clouds of MiTMAB-treated, LifeAct-mNG and mScar-Dyn2 co-expressing C2C12 cells at 48 hours post MyoD overexpression (**Supplementary Video 18**). mScar-Dyn2 (red arrowheads; 6-12 min) was observed highly enriched in the persistent actin cloud (green arrowheads). *n* = 3 independent experiments were performed with similar results. **f**, MHC immunostaining of Ctrl and dynamin1/2/3 tKO MEFs expressing wild-type Shi or Shi^4RD^ at five days post MyoD overexpression. **g**, Quantification of fusion index of the cells shown in (**f**). *n* = 3 independent experiments were performed (**Statistics Source Data-Fig.6g**). **h**, Immunostaining with anti-VASP and phalloidin in the MEFs described in (**f**). The VASP^-^ and VASP^tip^ protrusions are indicated by red arrowheads and white arrows, respectively. **i**, Quantification of the percentage of VASP^-^ protrusions shown in (**h**). *n* = 60 cells per genotype from three independent experiments were quantified (**Statistics Source Data-Fig.6i**). (**a-c**,**e**,**h**) Single plane confocal images. (**d**) Max z-projection of 8-10 focal planes from the ventral plasma membrane (z-step size: 500 nm). (**f**) Epifluorescence images. Mean ± s.d. values are shown in the dot plot (**j**) and bar-dot plot (**h**), and significance was determined by two-tailed student’s t-test. Sale bars: 3 μm (**a-c**), 2 μm (**d**,**e**), 5 μm (**h**) and 200 μm (**f**).

### Dynamin is required for actin bundling during invasive protrusion formation

To investigate which factor(s) are involved in organizing branched actin filaments into bundles, we first examined the localization of dynamin, a potent multi-filament actin-bundling protein required for protrusion invasiveness at the *Drosophila* fusogenic synapse^17^. Live imaging of MyoD^OE^ cells co-expressing mScar-dynamin 2 (Dyn2) and LifeAct-mNG revealed the presence of Dyn2 in the actin clouds (Fig. 6b, 2-3 min and Supplementary Video 15). This was not surprising because the Dyn2 protein was distributed throughout the cytosol. However, at a later time point, Dyn2 was enriched at the periphery of the actin bundle, followed by bundle thickening (Fig. 6b, 4-5 min), suggesting that Dyn2 may be in the process of crosslinking newly polymerized branched actin filaments back to the main bundle. In addition, live imaging of mScar-Dyn2 and Arp2-mNG co-expressing cells and immunostaining of fixed Arp2-mNG-expressing cells with anti-Dyn2 revealed that (Fig. 6c, 2-3 min and Supplementary Video 16) Dyn2 largely colocalized with Arp2-mNG within the actin clouds (Extended Data Fig. 7a-d). These observations suggest that Dyn2 may organize nascent branched actin filaments into bundles during protrusion growth.

To further test whether Dyn2 functions as a dynamic actin-bundling protein during invasive protrusion formation, we used a cell-permeable dynamin inhibitor, MiTMAB, known to inhibit dynamin GTPase activity by targeting the PH domain^60^. Although the PH domain is not required for dynamin helix-mediated actin bundling, it is indispensable for GTP-induced disassembly of full dynamin helices^17^. Failure in dynamin helix disassembly would “freeze” fully assembled dynamin-actin bundles and decrease the pool of cytosolic dynamin available to bundle new actin filaments^17^. Indeed, in the presence of MiTMAB, the dynamin-actin bundles formed normally, but dynamin helices failed to disassemble in the presence of GTP *in vitro* (Extended Data Fig. 7e), thus inhibiting the GTP-induced dynamic dynamin-actin bundling process. Correspondingly, treating MyoD^OE^ cells with MiTMAB that inhibits dynamin helix disassembly in the presence of cellular GTP resulted in persistent actin clouds that likely contain many “frozen” dynamin-actin bundles (Fig. 6d and Supplementary Video 17). Live imaging analysis confirmed persistent Dyn2 enrichment in these clouds (Fig. 6e; 6-12 min and Supplementary Video 18), in contrast to the lack of Dyn2 enrichment along the mature actin bundles in control cells (Fig. 6b; 6 min). Thus, by inhibiting dynamin helix disassembly, MiTMAB blocked the dynamic process of dynamin-mediated actin bundling and prevented the progression from actin clouds to mature actin bundles.

Next, we tested whether specifically disrupting the actin-bundling function of dynamin would impair actin bundle formation required for the generation of invasive protrusions. Our previous study identified four highly conserved, positively charged Arg (R) residues (R804, 829, 846 and 853) in the proline-rich domain of dynamin that are required for dynamin–mediated actin bundle formation^17^. Mutating these four Rs to negatively charged Asp (D) in *Drosophila* dynamin, Shibire (Shi), abolishes dynamin’s actin-bundling activity and renders the mutant protein (Shi^4RD^) non-functional in *Drosophila* S2R^+^ cell fusion^17^. To test the role for dynamin’s actin-bundling activity in invasive protrusion formation, we performed rescue experiments using wild-type Shi and Shi^4RD^ in dynamin-1/2/3 triple KO (tKO) mouse embryonic fibroblasts (MEFs)^61,62^ overexpressing MyoD. MyoD^OE^ transformed MEFs into MHC-expressing muscle cells (Fig. 6f), although the latter took a longer period of time (five days) to complete fusion than the MyoD^OE^ C2C12 cells (three days), consistent with the lower percentage of VASP^-^ invasive protrusions in MyoD^OE^ MEFs (8.7 ± 1.2%) compared to that in C2C12 cells (52.3 ± 6.1%). Dynamin tKO significantly impaired the fusion of MyoD^OE^ MEFs (Fig. 6f,g and Extended Data Fig. 7a), accompanied by a significant reduction in the number of invasive protrusions (1.0 ± 0.5%) (Fig. 6h,i), despite normal myogenic differentiation revealed by MHC expression (Fig. 6f). Expressing wild-type Shi in these cells rescued invasive protrusion formation (7.5 ± 0.5%) and cell-cell fusion (45.7 ± 5.8% compared to 62.8 ± 6.4% in wild type), whereas Shi^4RD^ expression failed to rescue protrusion formation (1.2 ± 0.9%) or cell-cell fusion (12.7 ± 4.4%) (Fig. 6f-i), suggesting that the actin-bundling activity of dynamin is required for invasive protrusion formation, the latter of which is indispensable for cell-cell fusion.

### Dynamin is aligned along nascent actin bundles in protrusions

To further characterize Dyn2’s role in actin bundling during protrusion initiation and growth, we performed immunofluorescence labeling of Dyn2 in detergent-extracted MyoD^OE^ cells (Fig. 7a-f and Extended Data Fig. 7f). Membrane dissolution by extraction exposes the cytoskeleton and its associated proteins^63^, such that they can be observed by various microscopic methods without the interference from membranes and/or cytosolic components. Immunofluorescence labeling of extracted cells revealed three distinct patterns of dynamin localization in the actin clouds – pattern 1: scattered Dyn2 punctae in the low actin density areas (Fig. 7a1z1; arrowheads); pattern 2: partially aligned Dyn2 punctae in the low actin density areas (Fig. 7a2z1; arrowheads); and pattern 3: well aligned Dyn2 punctae along actin bundles (Fig. 7a1z2, 7a2z2; arrows). Pattern 3 strongly supports Dyn2’s role in bundling actin, whereas pattern 2 may indicate an intermediate stage in the bundling process. Interestingly, few Dyn2 punctae were present in mature bundles with high F-actin content (Fig. 7a2z2; arrowheads), suggesting that Dyn2 is not required for stabilizing the mature bundles.

**Figure 7.**
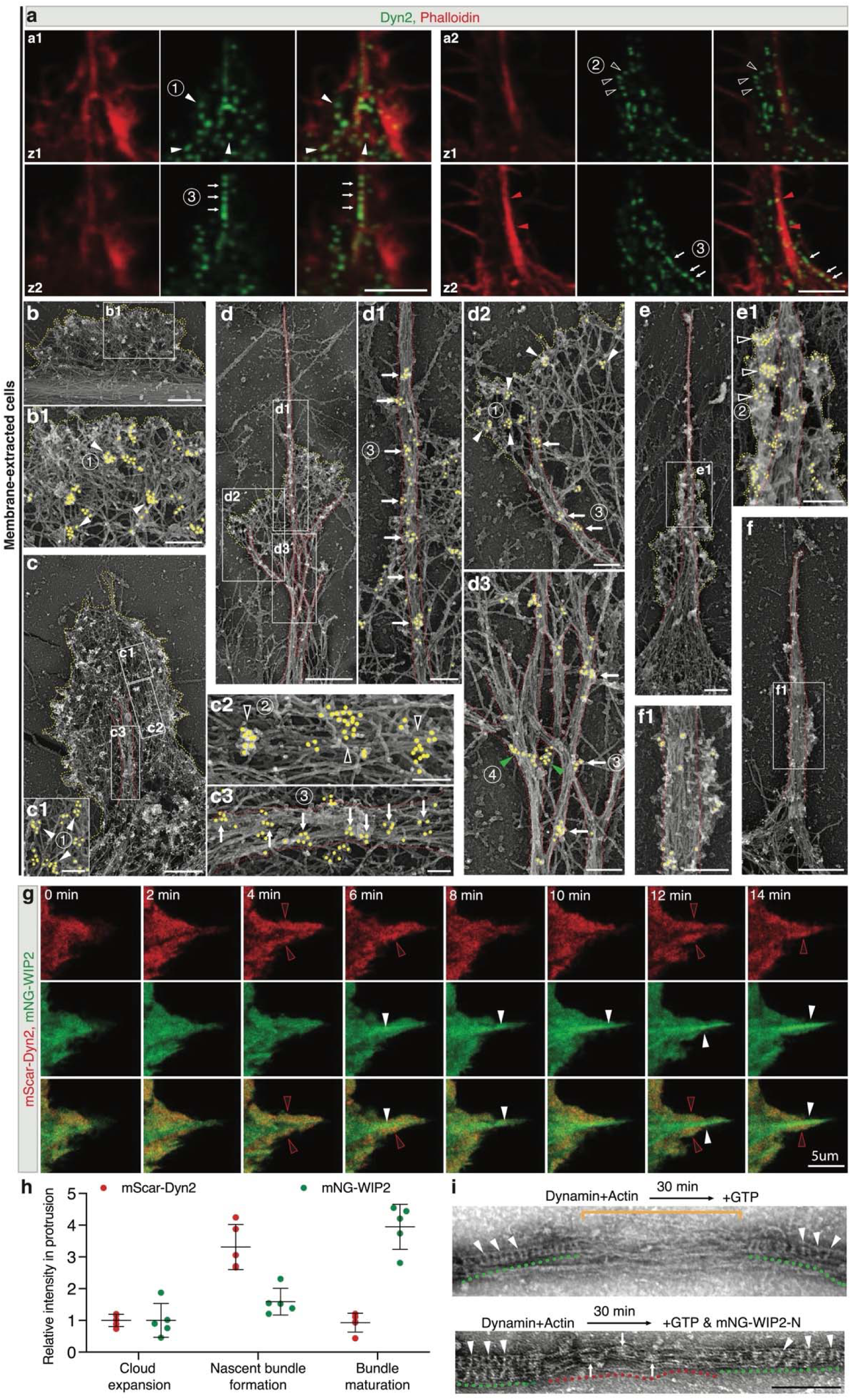
Dynamin is a “pioneer” actin-bundling protein for branched actin filaments and WIP2 stabilizes the actin bundles. **a**, Immunofluorescence labeling of Dyn2 in detergent-extracted MyoD^OE^ cells. Two (**a1**, **a2**) examples of actin clouds are shown, each with two consecutive focal planes (500 nm apart) from the ventral plasma membrane (bottom planes: a1z2 and a2z1; top planes: a1z1 and a2z2). Note that Dyn2 was localized in three patterns– scattered punctae in low-actin density areas (1; arrowheads in a1z1), partially aligned Dyn2 punctae in low-density area (2; hollow arrowheads in a2z1), or well aligned Dyn2 punctae along actin bundles (3; arrows in a1z2 and a2z2). In mature bundles with high F-actin content, only a few Dyn2 punctae were aligned with bundle (red arrowheads in a2z2). **b-f**, Dyn2 localization at ultrastructure level revealed by immunogold labeling and PREM of detergent-extracted cells. Dyn2 was localized in three patterns in actin clouds – 1 scattered clusters on branched actin filaments (white arrowheads in **b1**,**c1 and d2**), 2 partially aligned clusters on loosely organized actin filaments (hollow arrowheads in **c2** and **e1**), 3 clusters well aligned along nascent actin bundles (arrows in **c3**,**d1**,**d2** and **d3**). In addition, PREM revealed pattern 4 – clusters encompassing multiple bundles (green arrowheads in **d3**). In mature bundles with high F-actin content, a few small Dyn2 clusters were observed along thick bundle (**f** and **f1**). The boundaries of actin bundles and clouds are delineated by red and yellow dotted lines, respectively, in all panels. *n* ≥ 15 actin clouds for each stage were analyzed with similar results. **g**, Still images of live imaging of MyoD^OE^ cells co-expressing mNG-WIP2 and mScar-Dyn2 (**Supplementary Video 19**). mScar-Dyn2 and mNGT-WIP2 enrichment are indicated by red hollow and white arrowheads, respectively. **h**, Quantification of the relative intensity of mScar-Dyn2 and mNG-WIP2 in the protrusion during the indicated stages of protrusion formation. For each protein, the mean intensity in the entire protrusion at each stage was normalized by that at the cloud expansion stage. Mean ± s.d. values are shown in the dot plot (**Statistics Source Data-Fig.8h**). **i**, Electron micrographs of negatively staining of partially GTP hydrolysis-dissembled dynamin (Shi)-actin bundles treated without (top panel) or with (bottom panel) mNG-WIP-N. Rungs (several indicated by arrowheads) of dynamin helix are underlined by green dashed lines. The loosened actin bundle is indicated by a bracket (top panel), and mNG-WIP2-N patches on the tightened bundle (red dashed lines) are indicated by arrows (bottom panel). (**a**,**g**) Single planes confocal images. For (**a**,**g**,**h**), *n* = 3 independent experiments were performed with similar results. Scale bars: 3 μm (**a**), 120 nm (**c1**,**c2**),1 μm (**d**), 200 nm (**b1**,**d1**,**d2**,**d3**,**f1**,**i**), 100 nm (**c3**), 500 nm (**b**,**c**,**e**,**f**), 250 nm (**e1**) and 5 μm (**g**).

To observe the ultrastructural details of the Dyn2 punctae vs. the actin filaments, we performed immunogold labeling of Dyn2 using platinum replica electron microscopy (PREM) on membrane-extracted cells. Based on the size of the actin clouds and maturity of the actin bundles within the clouds, we have divided protrusion formation into five distinct stages – (**i**) actin cloud expansion, (**ii**) nascent bundle formation, (**iii**) proximal bundle crosslinking, (**iv**) distal bundle growth, and (**v**) bundle maturation (Fig. 7b-f). In stage (**i**), small actin clouds composed of branched, instead of linear, actin filaments were generated. Dyn2 was localized in scattered clusters on the branched actin filaments (Fig. 7b,b1; arrowhead, pattern 1), correlating with the scattered fluorescent Dyn2 punctae (Fig. 7a1z1; arrowheads, pattern 1) and the low-density Dyn2 in the actin clouds of intact cells (Fig. 6b, 2-3 min). In stage (**ii**), the actin cloud was larger and contained nascent actin bundles, along which Dyn2 clusters were aligned (Fig. 7c,c3; arrows, pattern 3), correlating with the fluorescent Dyn2 punctae on the actin bundles (Fig. 7a1z2, 7a2z2; arrows, pattern 3). In addition, scattered clusters of Dyn2 (Fig. 7c1; arrowheads, pattern 1) and partially aligned Dyn2 clusters on loosely organized actin filaments were observed (Fig. 7c2; arrowheads, pattern 2), the latter correlating with the partially aligned fluorescent Dyn2 punctae (Fig. 7a2z1; arrowheads; pattern 2). In stage (**iii**), more nascent bundles formed with Dyn2 clusters aligned along the bundles (Fig. 7d,d1,d2,d3; arrows, pattern 3). Moreover, nascent bundles were laterally connected at the proximal region of the protrusive structure (Fig. 7d3) with Dyn2 clusters encompassing these bundles transversely (Fig. 7d3; arrowheads; pattern 4), indicating a role for Dyn2 in crosslinking individual bundles together. In stage (**iv**), as the base of the protrusive structure became wider, small actin clouds could still be seen at the distal area of the actin bundle (Fig. 7e). Multiple Dyn2 clusters were partially aligned next to the main bundle (Fig. 7e1; arrowheads, pattern 2), correlating with the Dyn2 enrichment observed during protrusion growth in intact cells (Fig. 6b, 4 min; arrowheads). In stage (**v**), the mature protrusion contained densely packed actin filaments, the branching points of which were not visible anymore (Fig. 7f, f1), consistent with the decrease of Arp2 in the bundle (Fig. 6a, 6 min). Only a few small Dyn2 clusters were observed along the thick bundle (Fig. 7f1), correlating with the few fluorescent Dyn2 punctae on mature bundles (Fig. 7a2z2) and the decreased Dyn2 concentration along the mature protrusions in intact cells (Fig. 6b, 6 min). Taken together, our fluorescence imaging and ultrastructural analyses of extracted cells, as well as live imaging of intact cells, provide strong *in vivo* evidence supporting dynamin’s function as an actin-bundling protein that shapes the actin clouds into invasive protrusions.

### WIP2 stabilizes the dynamin-mediated actin bundles

Although dynamin is a highly efficient actin-bundling protein, fully assembled dynamin helices rapidly disassemble in the presence of GTP, leaving behind loosened actin bundles^17^. How the loose actin bundles are stabilized after the disassembly of dynamin helix remains unclear. Given WIP2’s ability to bundle actin (Fig. 5a-e and Supplementary Video 12,13), we asked whether WIP2 could be involved in bundle stabilization. As shown in Fig. 7g and Supplementary Video 19, live imaging of MyoD^OE^ cells expressing mScar-Dyn2 and mNG-WIP2 revealed a spatiotemporal coordination between Dyn2 and WIP2 during actin bundle formation. Dyn2 was initially present evenly in the actin cloud (2 min), followed by its enrichment at the periphery of the growing protrusion (4 min), and the subsequent convergence of enriched Dyn2 into the middle of the protrusion (6 min). This was when WIP2 became highly enriched in the actin bundle emerging from the center of the cloud (6 min), and remained enriched in the bundle even after Dyn2’s level decreased. These results support a model that dynamin is the “pioneer” actin-bundling protein that initiates the bundling process within the actin clouds. Once the dynamin full helices are disassembled by GTP hydrolysis, the loosely bundled actin filaments left behind would be crosslinked by WIP to form a tight and stable bundle. Indeed, negative-stain EM revealed that, upon GTP addition, stretches of loose actin bundles no longer associated with dynamin could be further crosslinked into tight bundles by WIP2 (Fig. 7h and Extended Data Fig. 7g).

### Branched actin polymerization promotes invasive protrusion formation and myoblast fusion during development

To examine whether branched actin polymerization is required for invasive protrusion formation and myoblast fusion *in vivo*, we knocked out ArpC2, a subunit of the Arp2/3 complex, to eliminate both N-WASP- and WAVE-mediated branched actin polymerization in the embryonic muscle progenitors using the MyoD^iCre^ mice, which starts to drive iCre expression at E9.75^64,65^. The mutant embryos (ArpC2^MyoD-KO^) exhibited normal gross morphology and body size at an early developmental stage (E12.5) but had curved back and thinner limbs at later stages (E15.5 and 17.5), indicating potential skeletal muscle developmental defects (Fig. 8a). Immunostaining with anti-MHC antibody showed significantly reduced limb muscle mass, cross-section area, and myofiber number in E17.5 mutant embryos (Fig. 8b-d and Extended Data Fig. 8e), whereas cell proliferation (marked by Ki67) and myoblast differentiation (marked by MyoG and MHC) remained unchanged (Extended Data Fig. 8a-d,f). Therefore, the small muscle size in ArpC2^MyoD-KO^ embryos was due to defects in myoblast fusion instead of in myoblast proliferation or myogenic/structural gene expression. In support of this, many differentiated (MHC^+^) but unfused, mononucleated myoblasts were seen attached to elongated myofibers in the longitudinal limb sections of E17.5 mutant embryos, compared to the few mononucleated myoblasts in control embryos at this stage (Fig. 8e,f). TEM analysis revealed that muscle cells in E15.5 ArpC2^MyoD-KO^ embryos seldom generated invasive protrusions at muscle cell contact sites (3.5% cells exhibited any protrusion; n = 57) compared to the control embryos (30% cells exhibited protrusions; n = 30) (Fig. 8g), suggesting that invasive protrusions propelled by branched actin polymerization are required for myoblast fusion *in vivo*. The presence of a small number of multinucleated myofibers in ArpC2^MyoD-KO^ mutant embryos (Fig. 8e) could be due to the presence of ArpC2 produced before MyoD iCre expression.

**Figure 8.**
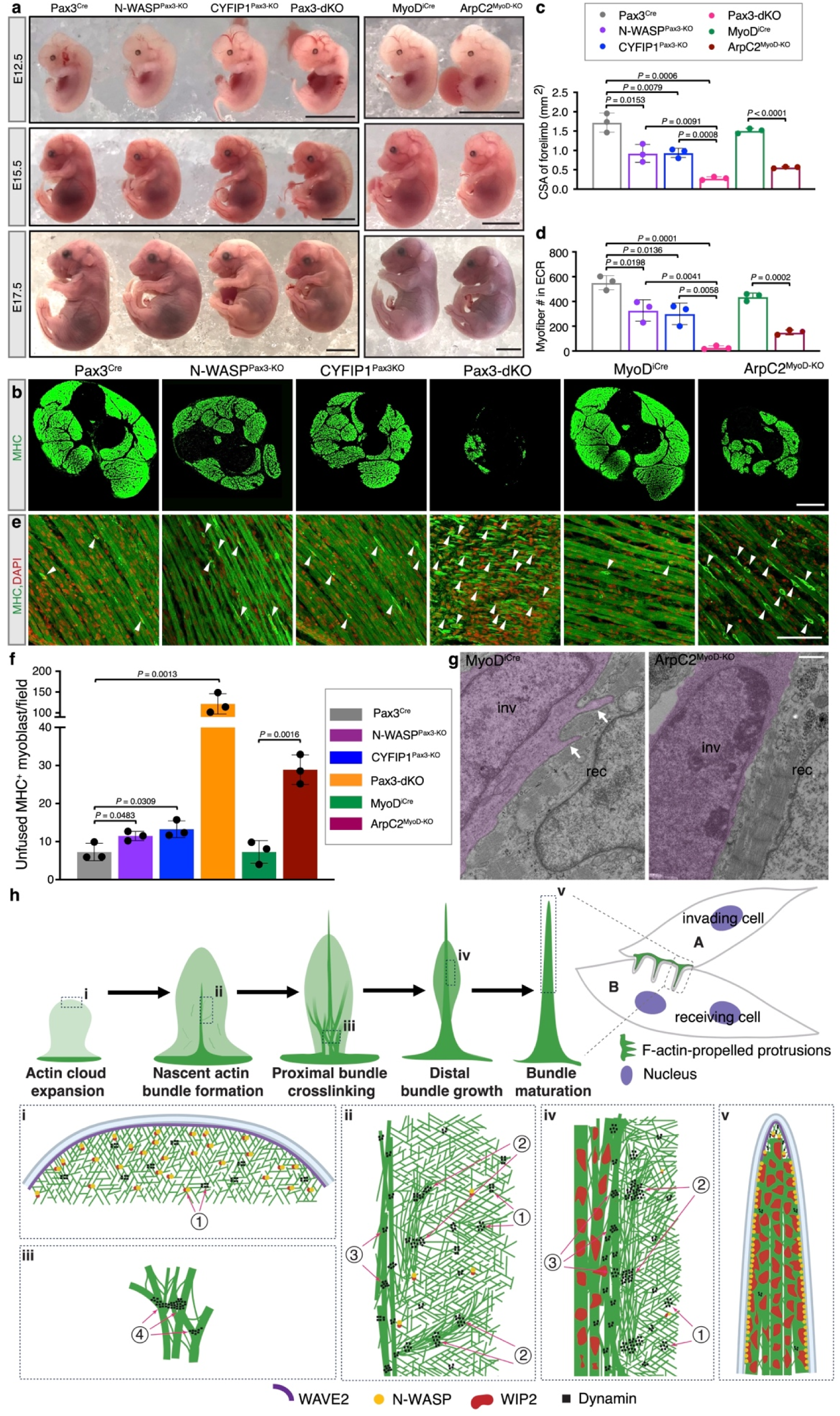
The Arp2/3 complex-mediated branched actin polymerization promotes invasive protrusion formation and fusion of mouse myoblasts *in vivo*. **a**, Gross morphology of control (Pax3^Cre^ or MyoD^iCre^ only), N-WASP^Pax3-KO^ (Pax3^Cre^; N-WASP^fl/fl^), CYFIP1^Pax3-KO^ (Pax3^Cre^; CYFIP1^fl/fl^), Pax3-dKO (Pax3^Cre^; N-WASP^fl/fl^; CYFIP1^fl/fl^), and ArpC2^MyoD-KO^ (MyoD^iCre^; ArpC2^fl/fl^) embryos at E12.5, E15.5, and E17.5, respectively. Note that the mutant embryos exhibited normal size and morphology at the early stage (E12.5), but abnormal spinal curvature and thinner limbs at late stages (E15.5 and E17.5). **b**, Immunostaining of forelimb cross section with anti-MHC of control and mutant embryos shown in (**a**) at E17.5. **c**, Quantification of the cross-section area (CSA) of the forelimb skeletal muscle in control and mutant embryos shown in (**c**). Three embryos of each genotype were examined (**Statistics Source Data-Fig.8c**). **d**, Quantification of the number of myofibers in the ECR muscle of forelimbs in control and mutant embryos shown in (**c**). Three embryos of each genotype were examined (**Statistics Source Data-Fig.8d**). **e**, Immunostaining for MHC in longitudinal sections of forelimbs in control and mutant embryos shown in (**b**). Randomly selected mononucleated MHC^+^ myoblasts are indicated by arrowheads. **f**, Quantification of the number of mononucleated MHC^+^ myoblasts shown in (**e**). Three embryos of each genotype were examined, respectively. Mononucleated MHC^+^ myoblasts in 12 40x microscopic fields of each embryo were counted (**Statistics Source Data-Fig.8f**). **g**, TEM analysis of the forelimb muscle of control and ArpC2^MyoD-KO^ embryos at E15.5. The invading fusion partner is pseudo-colored in magenta. The invasive protrusions are indicated by arrows. Nine out of 30 (30%) muscle cells in the control embryos and two out of 57 (3.5%) muscle cells in the ArpC2^MyoD-KO^ embryos were observed having invasive protrusions. *n* = 3 mice in each genotype were examined. **h-i**, A model describing the mechanisms of how invasive protrusions are generated from branched actin filaments at the mammalian fusogenic synapse. (**h**) Schematic diagrams of the five distinct stages of actin cytoskeletal rearrangements leading to invasive protrusion formation – (i) actin cloud expansion, (ii) nascent bundle formation, (iii) proximal bundle crosslinking, (iv) distal bundle growth, and (v) bundle maturation. (**i**) Schematic diagram of the actin polymerization and bundling process leading to the formation of an invasive protrusion. In stage (i), WAVE2 and N-WASP activate the Arp2/3 complex to generate a small lamellipodium-like structure (actin cloud) composed of branched actin filaments. WAVE2 is enriched at the leading edge, whereas N-WASP/WIP2 are broadly distributed in the cloud. Dynamin is distributed in scattered clusters (pattern 1) on the branched actin filaments. In stage (**ii**), the actin cloud is larger and, in some areas, the branched actin network has been organized into loosely aligned filaments or nascent bundles. Dynamin clusters are partially aligned on the loosely aligned actin filaments (pattern 2), or aligned along the nascent actin bundles (pattern 3). In stage (**iii**), more nascent bundles have formed in the proximal region of the protrusive structure, and some are laterally crosslinked to form a wider base. Dynamin clusters are seen encompassing the bundles, likely by engaging with common actin filaments in between (pattern 4). In stage (**iv**), small actin clouds continue to grow out from the distal part of the main bundle, with multiple dynamin clusters partially aligned next to it (pattern 2), contributing to bundle growth. Meanwhile, WIP has become highly enriched in the main bundle to stabilize it, with high level of N-WASP associated with the plasma membrane along the bundle. In stage (**v**), the mature protrusion contained densely packed actin filaments, the branching points of which were not visible anymore due to the tightly crosslinked filaments. Only a few small dynamin clusters are along the thick bundle, whereas more WIP is present to further tighten and stabilize the mature bundle. (**b**) Single planes confocal images. (**e**) max z-projection of 3 focal planes (z-step size: 1 μm). Mean ± s.d. values are shown in the bar-dot plot (**c**,**d**,**f**), and significance was determined by two-tailed student’s t-test. Scale bars: 5 mm (**a**), 0.5 mm (**b**), 100 μm (**e**) and 1 μm (**g**).

To further reduce the Arp2/3 complex-mediated actin polymerization during development, we used the Pax3^Cre^ mice, which starts to drive Cre expression in muscle progenitors at E8.5^66–68^. Since Pax3 and ArpC2 are localized on the same chromosome (which makes the generation of ArpC2^Pax3-KO^ technically challenging), we generated single conditional KO of N-WASP (N-WASP^Pax3-KO^) or CYFIP1 (a subunit of the WAVE complex) (CYFIP1^Pax3-KO^), as well as double conditional KO of N-WASP and CYFIP1 (Pax3-dKO) mice (Fig. 8a). While myoblast proliferation and myogenic/structural gene expression appeared normal in single and double conditional KO embryos (Extended Data Fig. 8a-d,f), the single KO mutants exhibited a moderate reduction of muscle mass at E17.5, which was exacerbated in Pax3-dKO mutant embryos (Fig. 8b-d and Extended Data Fig. 8e). In addition, the Pax3-dKO embryos showed a more severe myoblast fusion defect compared to the single mutants (Fig. 8e,f), suggesting that the N-WASP and WAVE proteins have partially redundant functions in promoting myoblast fusion during skeletal muscle development. Notably, Pax3-dKO embryos exhibited more significant muscle mass reduction and myoblast fusion defects than ArpC2^MyoD-KO^ embryos (Fig. 8b-f), consistent with Pax3^Cre^ being an earlier driver than MyoD^iCre^. Taken together, these results revealed a critical role for branched actin polymerization in promoting myoblast fusion *in vivo* and partially redundant functions for the N-WASP and WAVE family of actin NPFs in this process.

## Discussion

In this study, we identified the molecular components of the asymmetric fusogenic synapse in mammalian myoblast fusion and uncovered the mechanisms underlying the formation of branched actin-driven invasive protrusions. We found that two NPFs of the Arp2/3 complex, WAVE2 and N-WASP, exhibit largely distinct localization patterns in invasive protrusions. While WAVE2 is closely associated with the plasma membrane at the leading edge of the protrusive structures, N-WASP is enriched with WIP2 along the shafts of the protrusions. During protrusion initiation and growth, the Arp2/3 complex nucleates branched actin filaments to generate low density actin clouds, in which dynamin organizes new branched actin filaments into nascent bundles, followed by WIP2-mediated actin-bundle stabilization. Our findings revealed the spatiotemporal coordination of two actin NPFs (WAVE2 and N-WASP) and two actin-bundling proteins (Dyn2 and WIP2) in driving invasive protrusion formation at the fusogenic synapse (Fig. 8h,i).

Our previous work has demonstrated that invasive protrusions serve as a conserved mechanism underlying myoblast fusion in *Drosophila* and zebrafish, and this study has extended the conservation to mammals^14,20^. Interestingly though, the spatial organization of the invasive protrusions at the fusogenic synapse appears different between species. While the protrusions are clustered in both *Drosophila* and zebrafish, those at the mouse fusogenic synapses are more sparsely localized. This could be due to the different adhesion molecules involved in recruiting the actin polymerization machinery during myoblast fusion. In both *Drosophila* and zebrafish, Ig domain-containing cell adhesion molecules (CAMs) form a cluster at the fusogenic synapse to organize the invasive protrusions in a spatially restricted area^4,13^. It is unclear at present whether cell-cell adhesion plays a role in mammalian myoblast fusion, and, if so, which CAMs are involved. However, it has been shown that the cell-matrix adhesion molecule integrin is required for mammalian myoblast fusion^69,70^. Using a reconstituted cell-cell fusion system in *Drosophila*, we have demonstrated that integrin enhances cell-cell fusion by providing additional anchorage at the cell periphery, allowing the formation of numerous mechanically stiff, sparsely localized invasive protrusions along the cell-cell contact zone^18^. The large number of disperse protrusions along wide cell-cell contact zones would increase the probability of establishing fusogenic synapses between cells that are actively migrating, such as the mammalian muscle cells.

The sparsely localized invasive protrusions in mammalian myoblasts make it possible to visualize individual protrusions at high resolution and address a longstanding question – why do cells use two, instead of one, actin NPFs to generate invasive protrusions? In this study, we revealed the distinct spatial localization of WAVE2 and N-WASP in individual protrusions. WAVE2 exhibits specific and close association with the plasma membrane at the leading edge, thus allowing the generation of branched actin filaments right beneath the cell membrane. These filaments, in turn, provide seeds for the more broadly localized N-WASP to activate branched actin polymerization across the low density actin clouds. What makes N-WASP more unique is its ability to bind and stabilize the actin-bundling protein WIP2, the latter of which is highly enriched in the actin bundles. The WIP2-bound N-WASP, in turn, provides a rich pool of the latter beneath the cell membrane along the actin bundles for generating additional actin clouds to grow the protrusion.

Why do cells use two actin-bundling proteins, dynamin and WIP, to organize branched actin filaments into mechanically stiff bundles? Compared to bundling linear actin filaments, organizing branched actin filaments into directional bundles is a more complex process. Two features of the large GTPase dynamin make it an ideal pioneer bundling protein for branched actin filaments. First, dynamin can bundle multiple actin filaments (12 or 16 at its full capacity) while forming a helical structure^17^, making it possible to efficiently align and crosslink multiple filaments along the same direction.

Second, dynamin can mediate interactions between single helical bundles to generate “super bundles”^17^, further enhancing the efficiency of directional actin bundling. Thus, dynamin generates order out of a disordered, branched actin network more rapidly than any of the two-filament actin crosslinkers. However, the full helical structures of dynamin with actin filaments recruited to the outer rim are rapidly disassembled by GTP hydrolysis^17^, leaving behind loosely aligned parallel actin filaments. This is when a two-filament actin crosslinker, such as WIP2, exerts its function to stabilize the directional bundles. Thus, branched actin filament bundling requires a “handover” from Dyn2 to WIP2, two actin crosslinkers with distinct properties. Our finding of WIP2 as an actin-bundling protein helps to explain the essential roles for WIPs, all of which contain two similar WH2 domains, in increasing the invasiveness of finger-like protrusions in various cellular contexts. For example, protrusions are bendy in *Drosophila sltr* mutant^14^ and mouse WIP2^-/-^ myoblasts (Fig. 4a4,g), and mechanically stiff actin cores are lacking in podosomes and invadopodia of WIP1^-/-^ osteoclasts and metastatic carcinoma cells, respectively^71,72^.

Our study has demonstrated that invasive protrusions are distinct from filopodia, even though both are actin-rich finger-like protrusions and both arise from the cortical dendritic network formed by WAVE and the Arp2/3 complex^58,73^ (this study). Unlike the thin filopodia, the growth/extension of the thicker invasive protrusions are driven by branched, instead of linear, actin filaments. In addition, invasive protrusions are WIP^+^ along the shaft and VASP^-^ at the tip, whereas filopodia are WIP^-^ along the shaft and VASP^+^ at the tip. The different molecular and physical properties of these two types of protrusions correlate with their distinct functions – invasive protrusions made from short branched filaments are mechanically stiff and used by cells to invade their neighbors or basement membrane^3,14^, whereas filopodia composed of long and bendy linear filaments are less stiff and function as antennae for cells to probe their environment^52^. Given that branched actin polymerization is required for many cellular processes beyond cell-cell fusion, such as podosome and invadopodia formation in cancer metastasis, leukocyte diapedesis, immunological synapse maturation, and uterine-vulval attachment^71,72,74–76^, we speculate that similar molecular mechanisms are used to generate invasive protrusions in these other cellular contexts as well.

## Acknowledgements

We thank Dr. Pietro De Camilli for providing the tamoxifen-inducible dynamin 1/2/3 triple KO mouse embryonic fibroblasts. We also thank the Peter O’Donnell Jr. Brain Institute NeuroMicroscopy Core for access to the Abberior STED Facility Line super-resolution microscope system, and Drs. Kate Luby-Phelps and Dorothy Mundy of the UT Southwestern Quantitative Light Microscopy Core for training and assistance with acquiring the TIRF, SIM, and STED images. This work was supported by: an NIH grant (R35GM136316) to E.H.C., and a Fellowship from the UT Southwestern Hamon Center for Regenerative Science and Medicine (CRSM) to K.H.L. D.P. is an HHMI investigator.

## Author contributions

T.W. initiated the project. Y.L. and E.H.C. designed the project. Y.L., T.W., B.R., P.P., B.L., C.Y., R.Z., Z.L., and C.Z. performed experiments. Y.L. and E.H.C. collaborated with D.G., C.H., S.S., and R.L. on the mouse genetic experiments; B.L. and D.P. on the biochemistry experiments; and C.Y. and T.S. on the PREM experiments. B.R. and E.H.C collaborated with K.L. and D.S. on the micropattern preparations for live cell imaging experiments. Y.L., B.R., and E.H.C. analyzed the data. Y.L. and E.H.C. made the figures and wrote the manuscript. All authors commented on the manuscript.

## Extended Data Figures

**Extended Data Figure 1.**
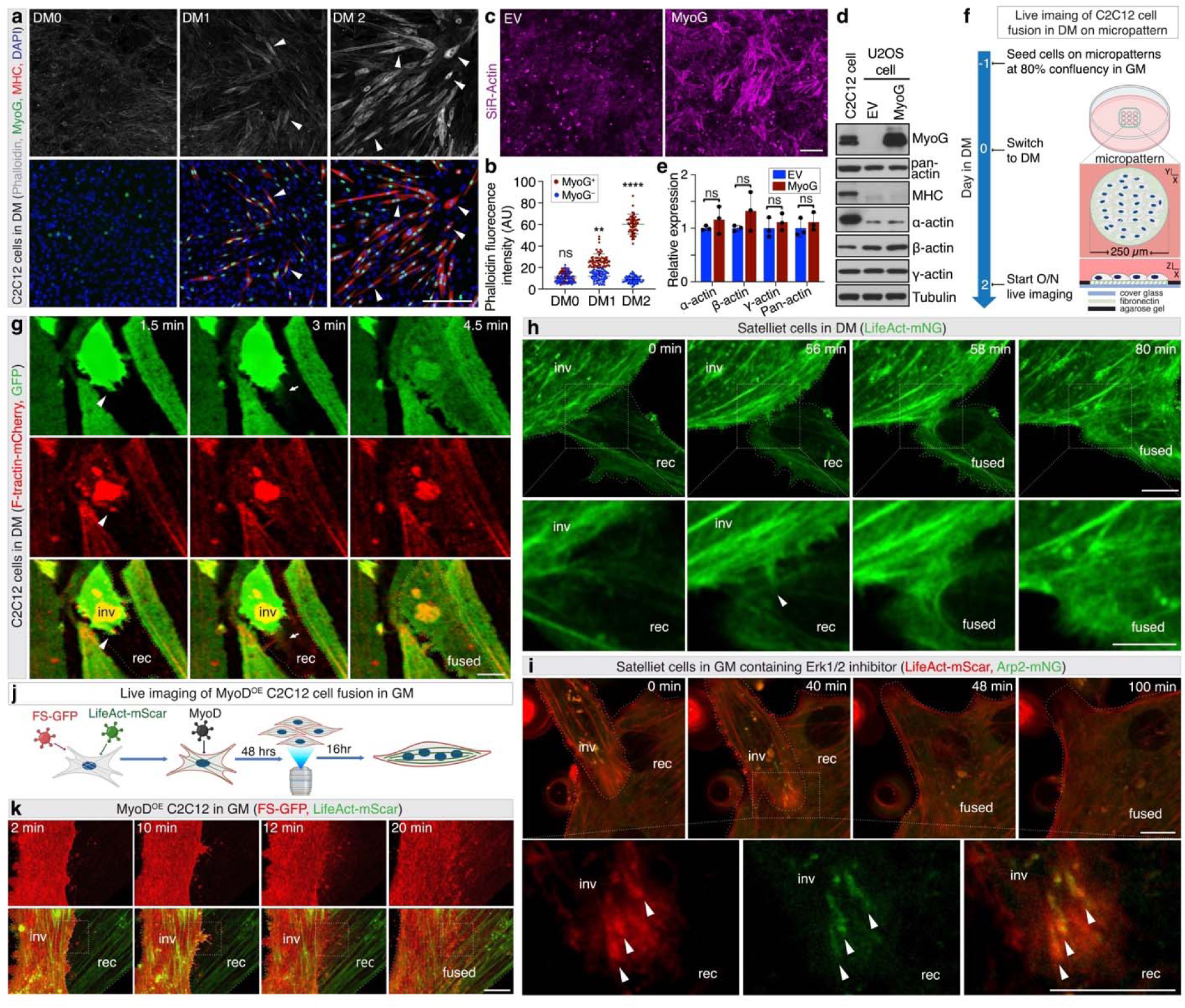
F-actin enriched invasive protrusions mediate the fusion of mouse C2C12 cells and satellite cells (a) The gradual increase in the F-actin level during mouse myoblast differentiation. Wild-type C2C12 cells cultured in differentiation medium (DM) were immunostained with anti-MyoG, anti-MHC, and phalloidin at the indicated time points (e.g. DM0 – day 0, DM1 – day 1, DM2 – day2). Note the gradual increase in the F-actin intensity in cells over time. A few randomly selected F-actin^high^ cells are indicated by arrowheads. (b) Quantification of the phalloidin signal intensity in cells shown in (**a**). MyoG^-^ or MyoG^+^ cells (n ≥ 18) at each indicated time point were measured for phalloidin intensity per experiment. *n* = 3 independent experiments were performed. Statistics source data can be found in “**Statistics Source Data-Extended Data Fig.1b**”. (c) MyoG expression increases F-actin level in U2OS cells. 80% confluent U2OS cells were infected by either empty lentiviral vector (EV) or lentivirus containing MyoG. Three days post-infection, the cells were incubated with SiR-Actin in culture medium for 30 minutes and fixed with 4% PFA. Note the increase in the F-actin level (indicated by the SiR-Actin intensity) in MyoG^OE^ cells (bottom panel) compared with the negative control (top panel). *n* = 3 independent experiments were performed with similar results. (d) MyoG expression does not change the levels of total actin protein or induce MHC expression in U2OS cells. U2OS cells shown in (**c**) were collected and subjected to WB with anti-α-, β-, γ-, pan-actin, anti-MyoG, and anti-MHC at day three post MyoG expression. C2C12 cells at day three in DM was used as a positive control for MyoG and MHC expression. Note that none of the actin isoforms or the total actin level was affected by MyoG expression. MHC was not induced in U2OS cells expressing MyoG. (e) Quantification of protein expression as shown in (**d**). The band intensity of each protein was normalized against β-tubulin. For each protein, the y axis indicates the expression in MyoG^OE^ cells relative to that in the EV cells. *n* = 3 independent experiments were performed. Statistics source data can be found in “**Statistics Source Data-Extended Data Fig.1e**”. (f) Schematic diagrams of live imaging C2C12 cell fusion in DM. C2C12 cells were first seeded at 80% confluency on micropatterns in growth medium (GM). Upon 100% confluency, the cells were cultured in DM for 48 hours and subjected to live imaging. (g) Still images from a time-lapse movie of a fusion event between F-tractin-mCherry-expressing C2C12 cells in DM. The F-tractin-mCherry^-^expressing C2C12 cells with or without cytosolic GFP were co-cultured, switched to DM, and subjected to live imaging as described in (**c**). Note that the F-actin-enriched membrane protrusion (1.5 min, arrowhead) led to fusion pore formation and GFP transfer from the top to the bottom cell (3 min, arrow) (see **Supplemental Video 3**). *n* = 23 fusion events were observed with similar results. (h) Still images from a time-lapse movie of a fusion event between two satellite cells cultured in DM. Low passage (≤ three passages) satellite cells were plated at 50% confluency on collagen type I-coated 6-well plates in satellite cell growth medium (SCGM) for 24 hours. The following day, the cells were infected by retrovirus containing LifeAct-mNG in SCGM. After 48 hours, the cells were trypsinized and plated at 40% confluence on fibronectin-coated cover glass in DM. After another 48 hours, the cells were subjected to live imaging. Note the presence of a prominent actin-enriched finger-like protrusion (56 min, arrowhead) prior to cell-cell fusion and LifeAct-mNG transfer from the invading cell to the receiving cell (58 min) (see **Supplemental Video 4**). *n* = 18 fusion events were observed with similar results. (i) Still images from a time-lapse movie of a fusion event between satellite cells cultured in GM with an Erk1/2 inhibitor. Low passage (≤ three passages) satellite cells were infected by a mixture of retroviruses containing LifeAct-mScar and Arp2-mNG in SCGM as described in (**g**). After 48 hours, the cells were trypsinized and plated at 40% confluence on fibronectin-coated cover glass in GM containing 1μM ERK1/2 inhibitor (SCH772984). After another 48 hours, the cells were subjected to live imaging. Note that the invading cell projected actin- and Arp2-enriched protrusions (40 min, arrowheads) prior to cell fusion (**Supplemental Video 5**). *n* = 7 fusion events were observed with similar results. (j) Schematic diagrams of live imaging MyoD^OE^ cell fusion in GM. C2C12 cells co-expressing LifeAct-mScar and FS-GFP were seeded at 30% confluency on fibronectin-coated cover glass in GM. After 24 hours, the cells were infected with retrovirus containing MyoD. 48 hours after MyoD overexpression, the culture medium was replaced with fresh GM and the cells were subjected to live imaging for 16 hours. (k) Zoomed-out view of the fusion event shown in Fig. 1g. The boxed areas are enlarged in Fig. 1g. (**a**,**c**,**i**) Max z-projection of 8-10 focal planes from the ventral plasma membrane (z-step size: 500nm). (**g**,**h**,**k**) Single plane confocal images. Mean ± s.d. values are shown in the line graph (**b**) and bar-dot plot (**e**), and significance was determined by two-tailed student’s t-test. **: *p* < 0.01, ****: *p* < 0.0001, ns: not significant. In (**g**,**h**,**i**,**k**), the cell boundaries are delineated by dotted lines. Scale bars: 50 μm (**a**), 200 μm (**c**), 10 μm (**h**, top panel); 5 μm (**h**, bottom panel) and 10 μm (**g**,**i**,**k**).

**Extended Data Figure 2.**
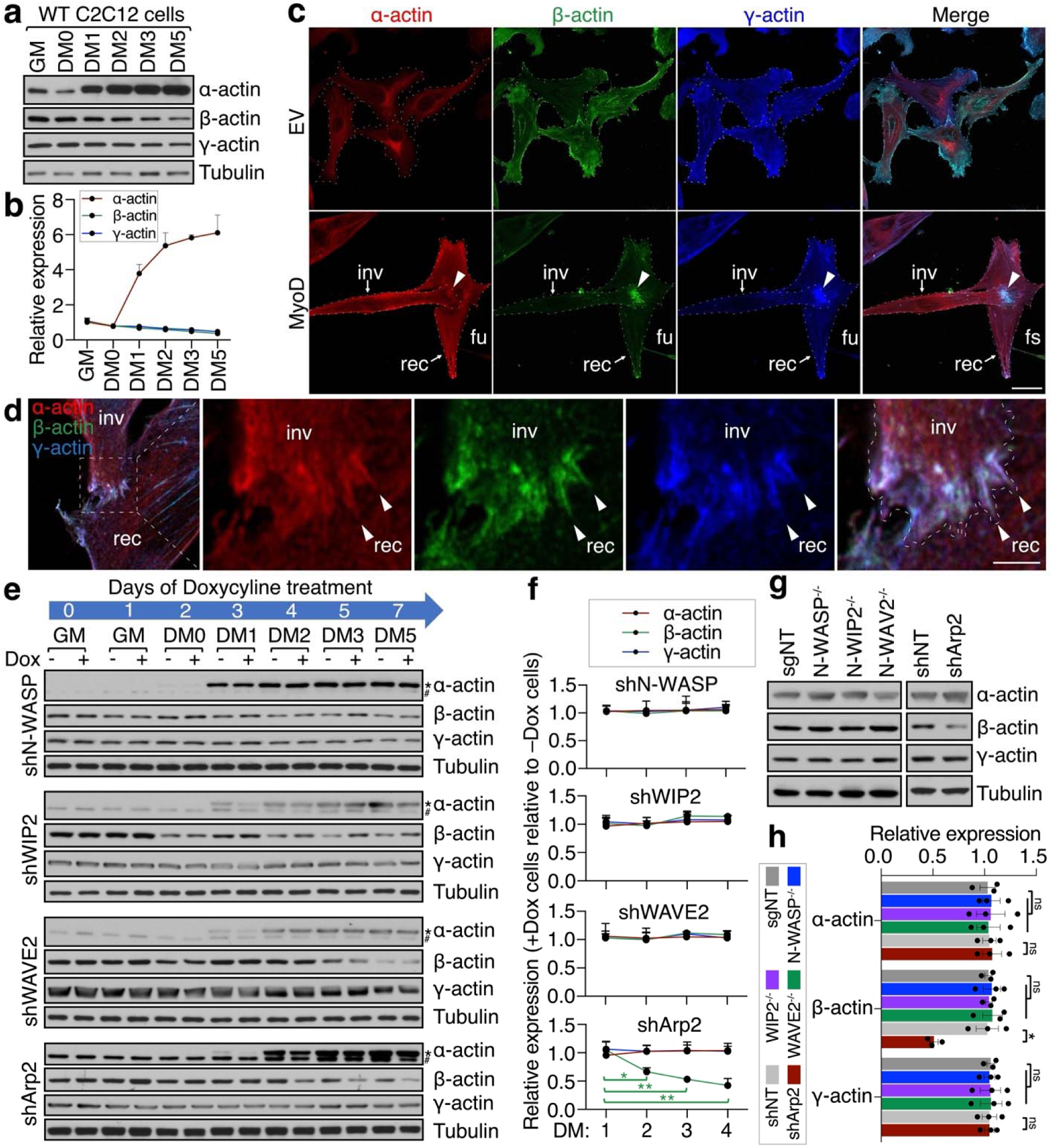
The expression and localization of sarcomeric and cytosolic actin isoforms during mouse myoblast fusion. **(a)** The sarcomeric (α-), but not the cytosolic (β- and γ-), actin level increases during C2C12 cell differentiation. Wild-type C2C12 cells were grown to 100% confluency in GM and then switched to DM. The cells were collected at the indicated time points during differentiation and subjected to western blot (WB) using antibodies specific for α-, β- and γ-actin, respectively. Note that α-actin expression significantly increased during differentiation, whereas the level of β- and γ-actin remained largely unchanged. **(b)** Quantification of protein expression shown in (**a**). The band intensity of each protein at each time point was normalized against β-tubulin. For each protein, the y axis indicates the expression relative to that in GM at the indicated time points. *n* = 3 independent experiments were performed. Statistics source data can be found in “**Statistics Source Data-Extended Data Fig.2b**”. **(c**,**d)** The invasive structure at fusogenic synapse contains all α-, β-, and γ-actin isoforms. Wild-type C2C12 cells were seeded at 60% confluence in GM. After 24 hours, the cells were infected by either empty retroviral vector (EV) or retrovirus containing MyoD. 48 hours post infection, the cells were fixed and subjected to immunostaining with anti-α-, β-, and γ-actin antibodies. The cell boundaries are delineated by dotted lines. Note that the fusogenic synapse (fs; arrowheads in **c**) contained all three actin isoforms with cytosolic β- and γ-actin highly enriched, and that α-actin exhibited an overall high level in the cells (**c**). Within the fusogenic synapse, all three actin isoforms were enriched in the invasive protrusions (arrowheads in **d**). *n* = 3 independent experiments were performed for each panel with similar results. **(e)** WB for α-, β-, and γ-actin in the doxycycline-induced conditional KD cells. Control (-Dox) and KD (+Dox) cells as described in Fig. 2a**,b** were harvested at the indicated time points to examine the α-, β-, and γ-actin levels by WB. **(f)** Quantification of protein expression as shown in (**e**). The band intensity of each protein at each time point was normalized against β-tubulin. For each protein, the y axis indicates its expression in KD relative to control cells. *n* = 3 independent experiments were performed. Statistics source data can be found in “**Statistics Source Data-Extended Data Fig.2f**”. **(g)** WB for α-, β-, and γ-actin in the N-WASP^-/-^, WIP2^-/-^, WAVE2^-/-^ KO, and Arp2 KD MyoD^OE^ cells. The control (expressing a non-targeting sgRNA (sgNT) or a non-targeting shRNA (shNT), respectively), N-WASP^-/-^, WIP2^-/-^, WAVE2^-/-^ KO, and Arp2 KD cells as described in Fig. 2e were harvested at 48 hours post MyoD overexpression to examine the protein levels of α-, β- and γ-actin by WB. **(h)** The quantification of protein expression shown in (**g**). The band intensity of each protein was normalized against β-tubulin. For each protein, the y axis indicates the expression of each group relative to the control cells. *n* = 3 independent experiments were performed. Statistics source data can be found in “**Statistics Source Data-Extended Data Fig.2h**”. Mean ± s.d. values are shown in the line graph (**b**,**f**), and significance was determined by two-tailed student’s t-test. Three independent experiments were performed. *: *p* < 0.05, **: *p* < 0.01, ns: not significant. (**c**) Max z-projection of 8-10 focal planes from the ventral plasma membrane (z-step size: 500nm). (**d**) Single plane confocal images. Scale bars: 30 μm (**c**) and 3 μm (**d**).

**Extended Data Figure 3.**
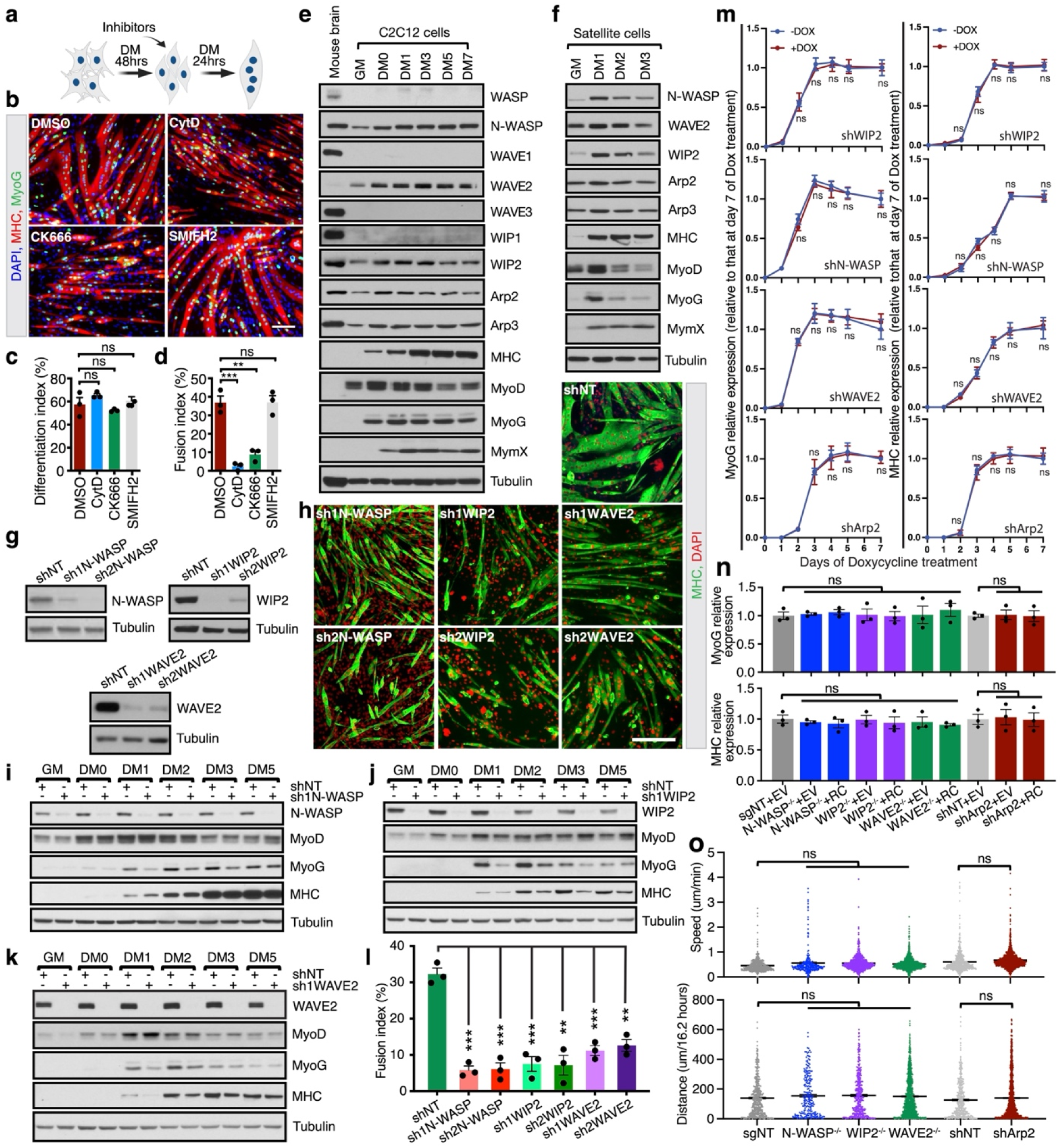
Branched actin polymerization promotes mouse myoblast fusion. **(a)** Schematic diagram of pharmacological treatment of C2C12 cells with actin polymerization inhibitors. Wild-type C2C12 cells were grown to 100% confluency in GM and then switched to DM. After 48 hours, the DM was supplemented with DMSO (0.05%), a general actin polymerization inhibitor cytochalasin D (CytD; 18nM), Arp2/3 complex inhibitor (CK666; 50 μM) or formin inhibitor (SMIFH2; 10 μM) dissolved in DMSO for 24 hours. **(b)** Branched, but not linear, actin polymerization promotes C2C12 cell fusion. The DMSO or inhibitor treated cells shown in (**a**) were immunostained with anti-MyoG and anti-MHC at day three in DM (24 hours post inhibitor treatment). Note the severe fusion defects caused by CytD and CK666 treatment, but not by DMSO and SMIFH2. **(c-d)** Quantification of the cell differentiation index (% of nuclei in MHC^+^ cells *vs*. total nuclei) and cell fusion index (% of nuclei in MHC^+^ myotubes (≥ 3 nuclei) *vs*. total nuclei) of the cells shown in (**b**). *n* = 3 independent experiments were performed. Statistics source data can be found in “**Statistics Source Data-Extended Data Fig.3c,3d**”. **(e)** N-WASP, WAVE2 and WIP2 are highly expressed in C2C12 cells. Wild-type C2C12 cells were grown to 100% confluency in GM and then switched to DM. The cells were collected at the indicated time points during differentiation and subjected to WB for NPFs, Arp2/3 complex, muscle differentiation markers, and the muscle fusogenic protein myomixer (MymX). Adult mouse brain tissue lysate was used as a positive control for NPFs and Arp2/3 complex expression. Note the high expression levels of N-WASP, WIP2, WAVE2, Arp2, and Arp3 among the homologous proteins. *n* = 3 independent experiments were performed with similar results. **(f)** N-WASP, WAVE2 and WIP2 are highly expressed in mouse satellite cells. Mouse satellite cells were grown to 70% confluence in SCGM and switched to DM. The cells were collected at the indicated time points during differentiation and subjected to WB for NPFs, Arp2/3 complex, muscle differentiation markers, and the muscle fusogenic protein MymX. *n* = 3 independent experiments were performed with similar results. **(g)** The knockdown (KD) efficiency of different shRNAs against target genes. C2C12 cells were infected by lentivirus (Lenti-shRNA-mCherry-puromycin) carrying a non-targeting shRNA (shNT) or shRNAs against the genes indicated. Two days later, cells underwent puromycin selection for five days. Then, the viable cells were harvested for WB. Two shRNAs were used to knock down each gene. Note the significant KD level of each target gene by both shRNAs. *n* = 3 independent experiments were performed with similar results. **(h)** C2C12 cell fusion is significantly inhibited by knocking down N-WASP, WIP2 and WAVE2 for seven days before differentiation. C2C12 cells were treated with the shRNAs described in (**g**) in GM, and underwent puromycin selection for five days. Once the cells reached 100% confluence after one day, the GM was replaced with DM. At day five of differentiation, the cells were immunostained with anti-MHC. **(i**,**j**,**k)** WB of the NPFs and muscle differentiation markers of the control and NPF KD cells described in (**h**). The control (shNT) and NPF KD C212 cells were collected at the indicated time points and subjected to WB. Note that at DM1, DM2, and DM3, MyoG was expressed at a lower level in KD cells compared to the control cells. *n* = 3 independent experiments were performed with similar results. **(l)** Quantification of the fusion index of C2C12 cells shown in (**h**). *n* = 3 independent experiments were performed. Statistics source data can be found in “**Statistics Source Data-Extended Data Fig.3l**”. **(m)** Quantification of MyoG and MHC protein expression in the doxycycline-induced conditional KD cells shown in Fig. 2b. The band intensity of each protein at each time point was normalized against β-tubulin. For each protein, the y axis indicates the expression at each time point relative to that at day seven of doxycycline treatment. *n* = 3 independent experiments were performed. Note that MyoG and MHC proteins levels were not significantly affected throughout differentiation. Statistics source data can be found in “**Statistics Source Data-Extended Data Fig.3m**”. **(n)** Quantification of MyoG and MHC protein expression in N-WASP^-/-^, WIP2^-/-^, WAVE2^-/-^, and Arp2 KD cells shown in Fig. 2e. The band intensity of each protein was normalized against β-tubulin. For each protein, the y axis indicates the expression in each group relative to the control (sgNT or shNT) cells. *n* = 3 independent experiments were performed. Note that MyoG and MHC proteins levels were not significantly affected. RC: rescue construct of the corresponding KD gene. *n* = 3 independent experiments were performed. Statistics source data can be found in “**Statistics Source Data-Extended Data Fig.3n**”. **(o)** Quantification of cell motility in the control (sgNT or shNT), N-WASP^-/-^, WIP2^-/-^, WAVE2^-/-^, and Arp2 KD cells. The cytosolic mScar-expressing sgNT, N-WASP^-/-^, WIP2^-/-^, WAVE2^-/-^, shNT, and Arp2 KD cells were generated by retroviral infection. Infected cells were then plated in GM at 50% confluency. After 24 hours, the cells were subjected to live imaging for 16 hours (see **Supplemental Video 7**). Each cell was tracked by TrackMate using ImageJ and is shown as one dot in the dot plot. *n* ≥ 223 cells in each genotype from three independent experiments were analyzed. Statistics source data can be found in “**Statistics Source Data-Extended Data Fig.3o**”. (**b**,**h**) Epifluorescence images. Mean ± s.d. values are shown in the line graph (**m**), bar-dot plot (**c**,**d**,**l**,**n**) and dot plot (**o**), and significance was determined by two-tailed student’s t-test. **: *p* < 0.01, ***: *p* < 0.001, ns: not significant. Scale bars: 50 μm (**b**,**h**).

**Extended Data Figure 4.**
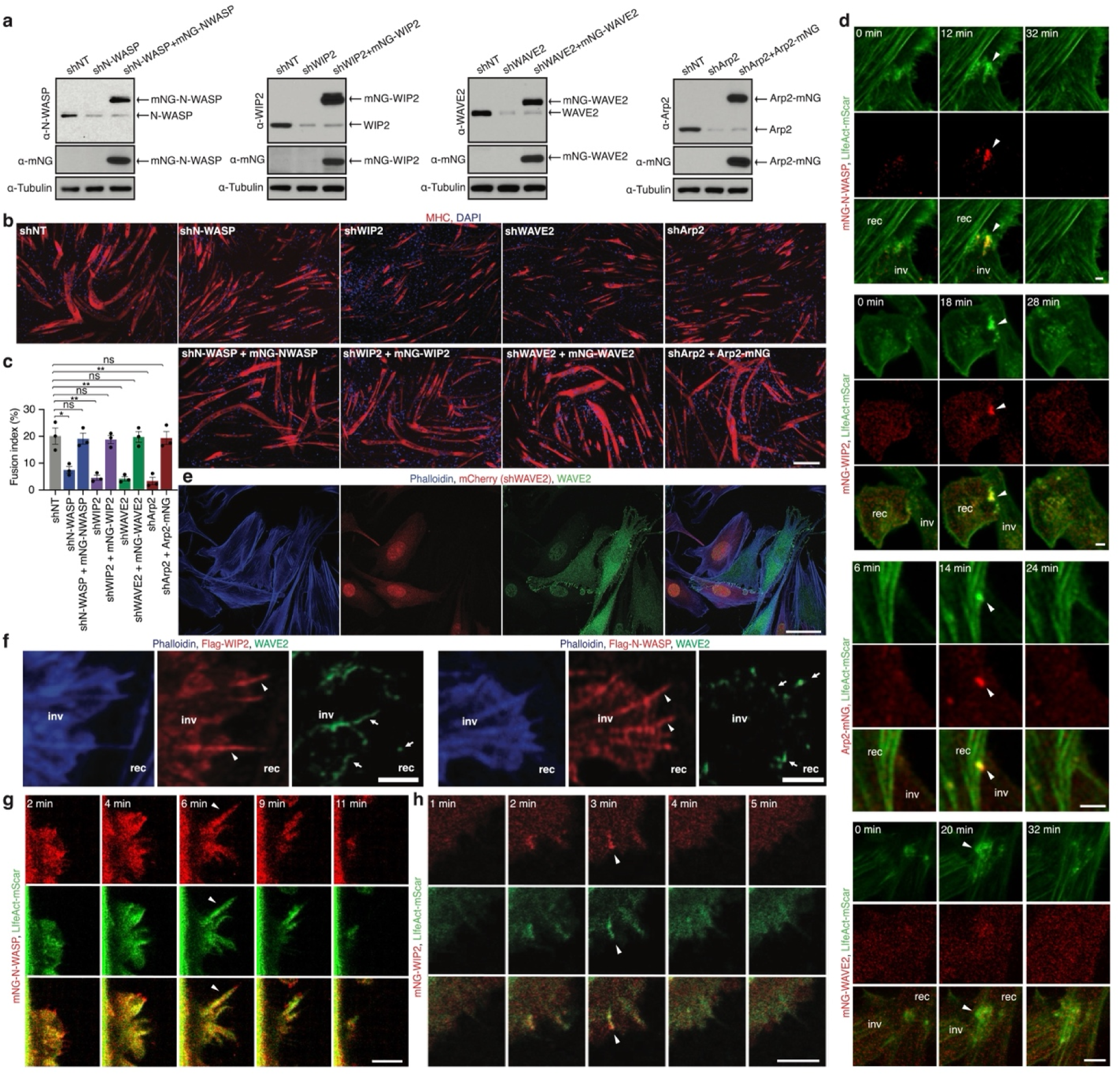
The dynamic enrichment of NPFs in the invasive protrusions at the fusogenic synapse. **(a,b)** mNG-tagged N-WASP, WIP2, WAVE2, and Arp2 rescue the fusion defects in N-WASP-, WIP2-, WAVE2- and Arp2-KD cells, respectively (see **Extended Data** Fig. 3g,h for the KD efficiency and effects). The KD cells were infected by retrovirus containing the corresponding KD gene with an mNG tag. After two days, the cells were plated and allowed to grow in GM into 100% confluence, before switching into DM. At day five in DM, the cells were collected for WB (**a**) and immunostaining (**b**). Note the expression of mNG-tagged proteins shown in (**a**) and the rescue of fusion shown in (**b**). **(c)** Quantification of the cell fusion index of each genotype shown in (**b**). *n* = 3 independent experiments were performed. Mean ± s.d. values are shown in the bar-dot plot, and significance was determined by two-tailed student’s t-test. *: *p* < 0.05, **: *p* < 0.01, ns: not significant. Statistics source data can be found in “**Statistics Source Data-Extended Data Fig.4c**”. **(d)** Single focal plane stills from a time-lapse movie of the fusion events shown in Fig. 3a. Note that the F-actin enrichments (arrowheads) at the fusogenic synapse are similar to those in Fig. 3a (see **Supplemental Video 8**). **(e)** Testing the specificity of the WAVE2 antibody used in this study. The WAVE2 KD C2C12 cells as shown in **Extended Data** Fig. 3g were mixed with wild-type C2C12 cells and seeded at 30% confluence in GM. After 24 hours, the cells were immunostained with anti-WAVE2 and phalloidin. The WAVE2 KD cells were marked by mCherry expression and showed little anti-WAVE2 staining (the nuclear signal was non-specific). *n* = 3 experiments were performed with similar results. **(f)** Single-channel images for the merged images shown in Fig. 3b**,c**. **(g,h)** Stills images from a time-lapse movie showing that N-WASP (**f**) and WIP2 (**g**) colocalized with F-actin along the shafts of straight protrusions. C2C12 cells co-expressing LifeAct-mScar and mNG-N-WASP (or mNG-WIP2) were infected with retrovirus containing MyoD. At 48 hours post MyoD overexpression, the cells were subjected to live imaging (see **Supplemental Video 9 and 10**). Single plane confocal images with F-actin enrichment are shown here. Arrowheads point to randomly selected protrusions. *n* = 3 experiments were performed with similar results. (**b**) Epifluorescence images. (**d,f,g,h**) Single plane confocal images. (**e**) Max z-projection of 8-10 focal planes from the ventral plasma membrane (z-step size: 500nm). Scale bars: 200 μm (**b**), 30 μm (**e**), 2 μm (**f**) and 5 μm (**d,g,h)**.

**Extended Data Figure 5.**
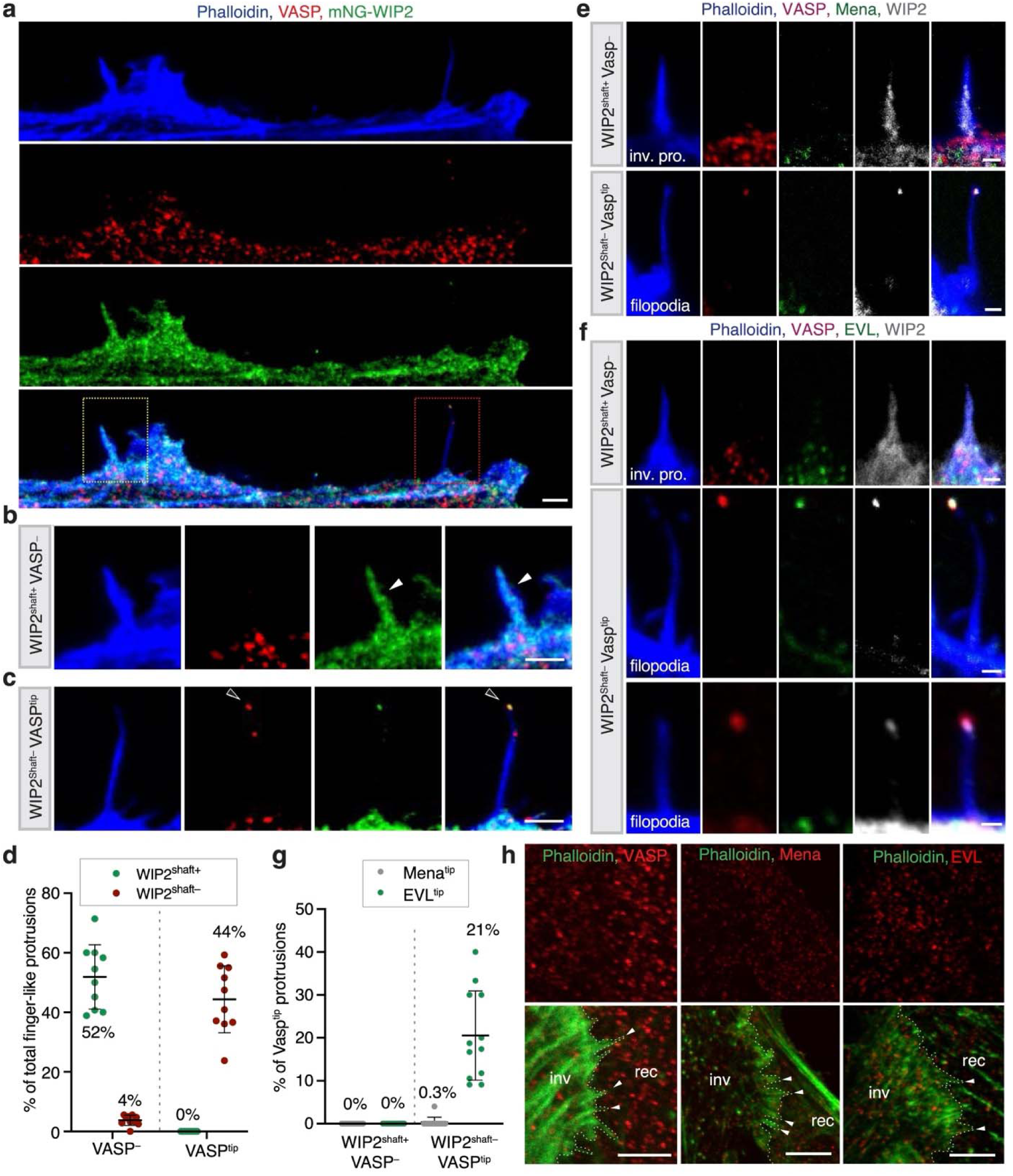
C2C12 cells generate both invasive protrusions and filopodia during differentiation. **(a)** Immunostaining with anti-VASP in the C2C12 cells expressing mNG-WIP2 at day two of MyoD overexpression. The cortical area of a randomly picked cell is shown. **(b)** Enlarged view of yellow-boxed area in **(a)**. Note that mNG-WIP2 was enriched in the shaft of the VASP^-^ protrusion (invasive). **(c)** Enlarged view of red-boxed area in **(a)**. Note that mNG-WIP2 was not enriched in the shaft of VASP^tip^ filopodia (non-invasive). **(d)** WIP2 is enriched in the shafts of VASP^-^, but not VASP^tip^, protrusions. The WIP2 enrichment was analyzed in the VASP^-^ and VASP^tip^ protrusions in mNG-WIP2-expressing cells at day two of MyoD overexpression. Note that the cells projected two major types of protrusions, ∼52% WIP2^shaft+^VASP^-^ and ∼44% WIP2^shaft–^VASP^tip^ protrusions. The rest of the protrusions (4%) were short and WIP2^shaft–^VASP^-^. *n* = 301 protrusions from ten randomly selected cells were analyzed. Mean ± s.d. values are shown in the dot plot. Statistics source data can be found in “**Statistics Source Data-Extended Data Fig.5d**”. (**e-g**) Mena and EVL are not enriched in the invasive protrusions. The mNG-WIP2-expressing cells was immunostained with anti-Mena or anti-EVL, together with anti-VASP and phalloidin at day two of MyoD overexpression. Randomly selected WIP2^shaft+^VASP^-^ invasive protrusions (inv. pro.) and WIP2^shaft–^VASP^tip^ filopodia are shown for Mena (**e**) and EVL (**f**) labeling. Quantification of the percentage of Mena^+^ and EVL^+^ protrusions is shown in (**g**). Note that neither Mena or EVL was localized in the invasive protrusions (0 out of *n* ≥ 90 WIP2^shaft+^VASP^-^ protrusions). EVL was co-enriched with VASP at the tips of ∼21% filopodia (41 out of *n* = 199 WIP2^shaft–^VASP^tip^ filopodia), and Mena was rarely observed at the tip of any filopodia (1 out of *n* = 186 WIP2^shaft–^ VASP^tip^ filopodia). Mean ± s.d. values are shown in the dot plot (**g**). Statistics source data for **g** can be found in “**Statistics Source Data-Extended Data Fig.5g**”. **(h)** Ena/VASP family members are not enriched in the invasive protrusions at the fusogenic synaspe. After day two of MyoD overexpression, wild-type C2C12 cells were immunostained with anti-VASP, anti-Mena, anti-EVL, and phalloidin. Note that none of VASP, Mena, or EVL was enriched at the tips of the invasive protrusions at the fusogenic synapse (0 out of *n* ≥ 10 fusogenic synapse for each protein). Cell boundaries are delineated by dotted lines. (**a-c,e-g,i**) Single plane confocal images. Scale bars: 2 μm (**a,b,c**), 3 μm (**e**), 1 μm (**f,g**) and 10 μm (**i**).

**Extended Data Figure 6.**
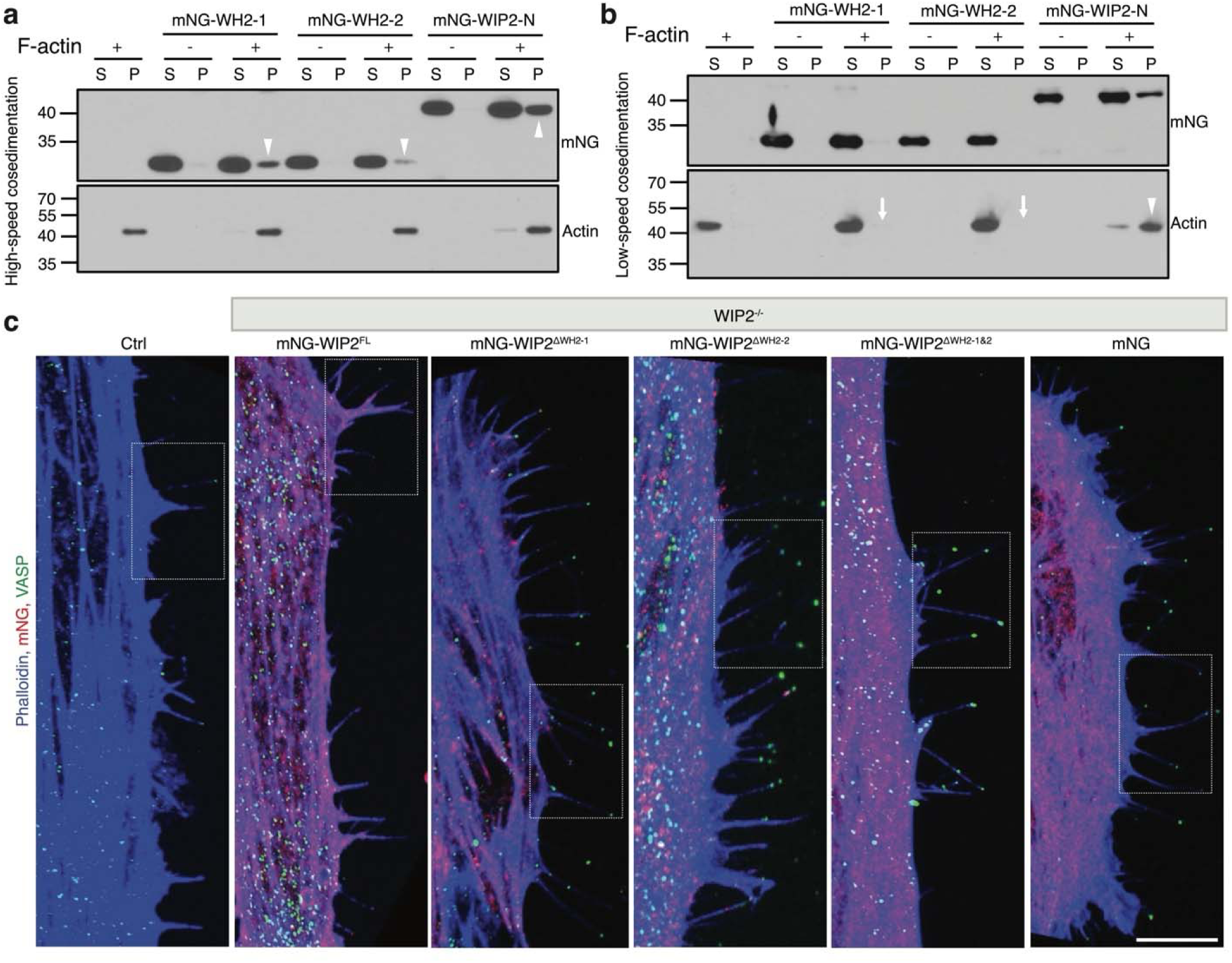
Both WH2 domains of WIP2 are required for actin bundling and invasive protrusion formation. **(a-b)** High-speed and low-speed F-actin co-sedimentation assays for the WH2 domains of WIP2. 4 μM mNG-WH2-1 (mNG-tagged first WH2 domain (aa 35-53)), mNG-WH2-2 (mNG-tagged second WH2 domain (aa 80-105)), or mNG-WIP2-N (mNG-tagged N-terminal domain of WIP2 containing both WH2-1 and WH2-2 (aa 1-176)) was incubated with or without 5 μM F-actin at room temperature (RT). After 30 minutes, the samples were subjected to high-speed (**a**; 100,000g) or low-speed (**b**; 13,600g) co-sedimentation assays. After centrifugation, the supernatant (S) and pellet (P) were subjected to SDS-PAGE. In (**a**), all three purified proteins bound F-actin (present in the pellet only when incubated with F-actin) (arrowheads), although mNG-WH2-2 showed relatively weaker binding compared to mNG-WH2-1. In (**b**), F-actin was bundled by mNG-WIP2-N (arrowhead), but not by mNG-WH2-1 or mNG-WH2-2 (arrows). *n* = 3 independent experiments were performed for (**a**) and (**b**) with similar results. **(c)** Zoomed-out view of the cells shown in Fig. 5f. The boxed areas are enlarged in Fig. 5f. A single plane confocal image is shown for each image. Scale bar: 10 μm.

**Extended Data Figure 7.**
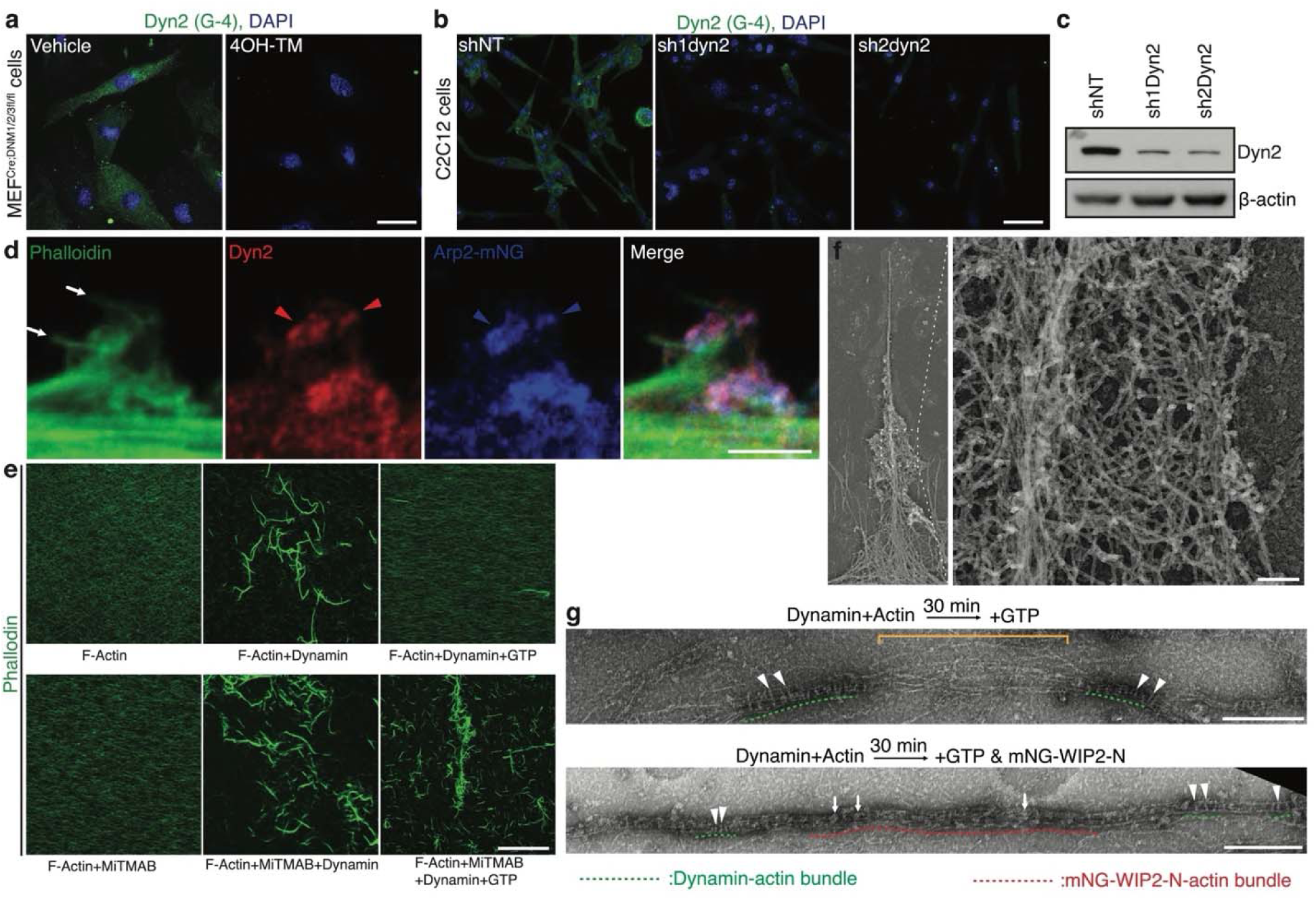
Both dynamin and WIP2 are involved in actin bundling during invasive protrusion formation. **(a-c)** The Dyn2 antibody (G-4) specifically recognizes Dyn2. (**a**) Tamoxifen-inducible dynamin 1/2/3 triple KO (tKO) MEFs were cultured with or without 2 μM 4-hydroxytamoxifen (4OH-TM) in the GM for 72 hours, and subsequently fixed for immunostaining with anti-Dyn2. (**b**) C2C12 cells were infected by lentivirus containing shRNAs against Dyn2. After 48 hours, the cells underwent puromycin selection for five days, and the viable cells were fixed for immunostaining (**b**) or harvested for WB (**c**) using anti-Dyn2. Note that anti-Dyn2 did not detect any signal in tKO cells and only greatly reduced signals in dyn2 KD cells, indicating its specificity. *n* = 3 independent experiments were performed with similar results. **(d)** Dyn2 and Arp2 are largely co-localized in the actin clouds. Arp2-mNG-expressing C2C12 cells were immunostained with anti-Dyn2 and phalloidin at day two of MyoD overexpression. Note the co-localization of Arp2 and Dyn2 within the actin clouds (arrowheads) at the periphery of actin bundles (arrows). *n* = 3 independent experiments were performed with similar results. **(e)** MiTMAB inhibits the disassembly of dynamin helices in the presence of GTP. F-actin pre-assembled from 1 μM G-actin was incubated with or without 1 μM dynamin (Shi) for 30 minutes at RT. Subsequently, 2mM GTP with or without 40 μM MiTMAB was added to the appropriate reaction mixtures and incubated for five minutes. The samples were then mixed with Alexa Fluor 488-conjugated phalloidin and subjected to confocal imaging on glass coverslips. Note that the dynamin-induced actin bundles were disassembled by GTP addition (top right panel). MiTMAB did not bundle actin by itself (bottom left panel) or inhibit dynamin-induced actin bundling (bottom middle panel), but it inhibited dynamin helix disassembly in the presence of GTP, thus leaving the actin bundles intact (bottom right panel). *n* = 3 independent experiments were performed with similar results. **(f)** Specificity test of the HA antibody used in the immunogold labeling experiments. Boxed area is enlarged on the right. Dyn2 KD C2C12 cells at day two of MyoD overexpression were stained with anti-HA for immunogold labeling, followed by PREM analysis. Note the absence of gold particles in these cells, demonstrating the specificity of the primary anti-HA antibody and the secondary antibody. *n* = 13 protrusions were imaged with similar results. **(g)** An additional example showing that WIP2 stabilizes actin bundles after dynamin helix disassembly. As described in Fig. 7h, dynamin-actin bundles were incubated with 1mM GTP without (top panel) or with (bottom panel) mNG-WIP2-N for 30 min, and subjected to negative-stain EM. Note that GTP addition resulted in partial disassembly of the dynamin helix. Residual rungs (several indicated by arrowheads) of the dynamin helix are underlined by green dashed lines. The loosened actin bundle is indicated by a bracket (top panel), and the mNG-WIP2-N patches (light-colored) on the tightened bundle (red dashed lines) are indicated by arrows (bottom panel). (**a,b**) Epifluorescence images. (**d,e**) Single plane confocal images. Scale bars: 50 μm (**a,b,e**), 4 μm (**d**), 100 nm (**f**) and 300 nm (**g**).

**Extended Data Figure 8.**
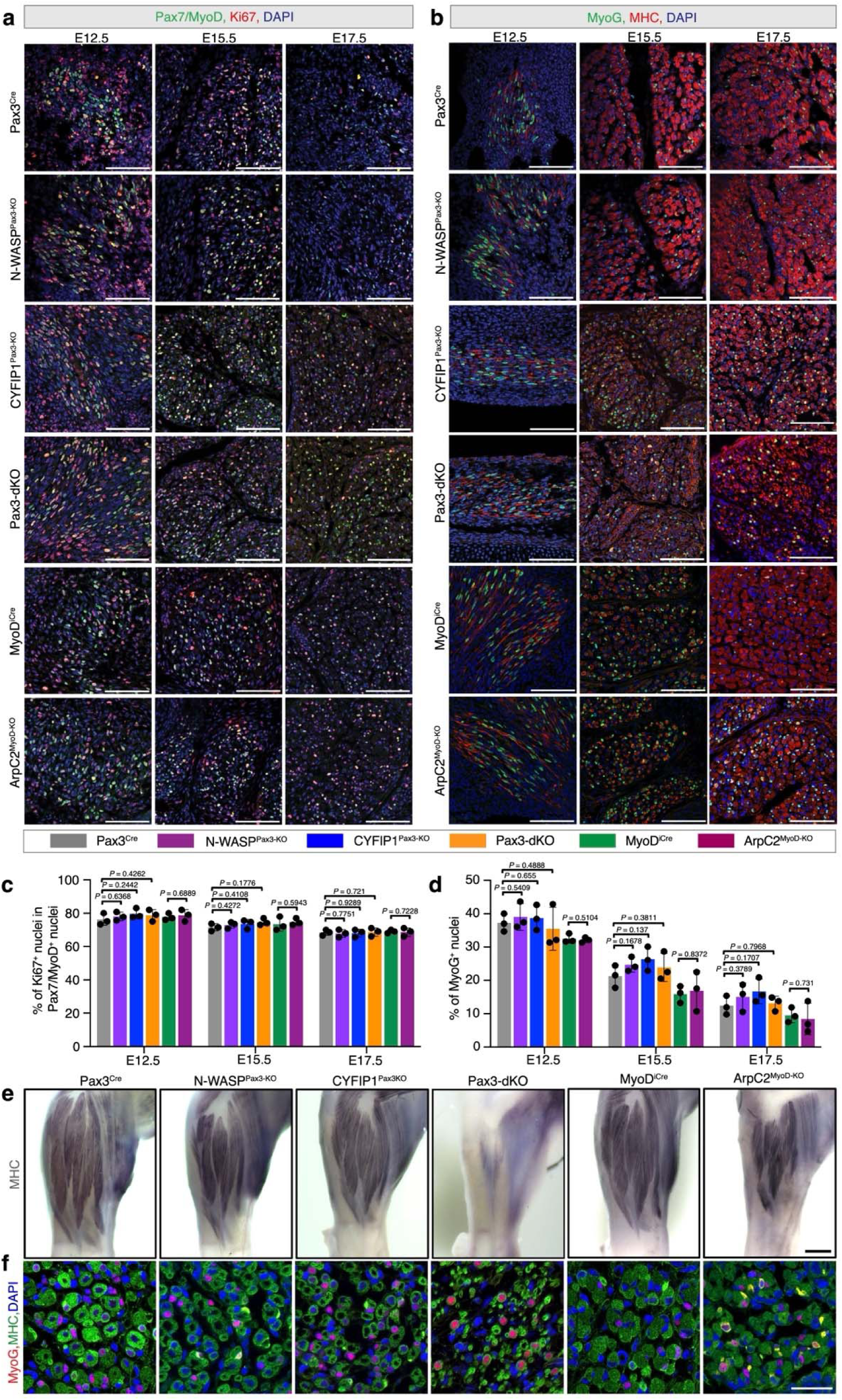
Branched actin polymerization is not required for myoblast proliferation and muscle-specific gene expression during skeletal muscle development. **a**, Immunostaining for Ki67 (proliferation maker), MyoD and Pax7 (muscle progenitor markers) in cross sections of the forelimb muscle of control (Pax3^Cre^ or MyoD^iCre^ only), N-WASP^Pax3-KO^, CYFIP1^Pax3-KO^, Pax3-dKO, and ArpC2^MyoD-KO^ embryos at E12.5, E15.5, and E17.5 as shown in Fig.8**. b**, Immunostaining for MyoG and MHC in cross sections of the forelimb muscle of the control and mutant embryos shown in (**a**) at E12.5, E15.5, and E17.5. **c**, Quantification of the proliferating myoblasts of the genotypes shown in (**a**). The percentage of Ki67^+^Pax7/MyoD^+^ nuclei in Pax7/MyoD^+^ nuclei was quantified. Note that Ki67 was normally expressed in the myoblast progenitors of the mutant embryos (**Statistics Source Data-Extended Data Fig.8c**). **d**, Quantification of the percentage of MyoG^+^ nuclei out of the total nuclei in the samples shown in (**b**). Note that MyoG was normally expressed in the muscle progenitors of the mutant embryo (**Statistics Source Data-Extended Data Fig.8d**). **e**, Whole mount immunostaining with anti-MHC of forelimbs of control and mutant embryos shown in (**Fig.8a**) at E17.5. *n* = 3 embryos of each genotype were examined with similar results. **f**, Immunostaining for MyoG and MHC in the forelimb cross sections of control and mutant embryos as shown in (**Fig.8b**) at E17.5. *n* = 3 embryos of each genotype were examined with similar results. For (**c,d**), *n* = 3 embryos of each genotype were examined for each time point, respectively. Myoblasts in 12 40x microscopic fields of each embryo were examined. Mean ± s.d. values are shown in the bar-dot plots, and significance was determined by two-tailed student’s t-test. (**a,b,e,f**) Single plane confocal images. Scale bars: 100 μm (**a,b**), 0.3 mm (**e**), and 100 μm (**f**).

## Supplementary Video Legends

**Supplementary Video 1.** Myoblast fusion occurs between C2C12 cells containing high level of F-actin Time-lapse imaging of C2C12 myoblast fusion at day two in differentiation medium (DM) on a micropattern. SiR-Actin was added to the medium for 30 minutes to label the cellular F-actin prior to live imaging. The time interval is ten minutes. Max z-projection of 8-10 focal planes from the ventral plasma membrane (z-step size: 500 nm) is shown. Scale bar: 100 μm.

**Supplementary Video 2.** Invasive protrusions mediate C2C12 cell fusion Time-lapse imaging of fusion between GFP^+^ and GFP^-^ C2C12 cells at day two in DM on a micropattern. Note the appearance of three invasive membrane protrusions (arrowheads) at the fusion site immediately before cell fusion, indicated by GFP transfer from the invading cell to the receiving cell. The time interval is two minutes. Max z-projection of 8-10 focal planes from the ventral plasma membrane (z-step size: 500 nm) is shown. Scale bar: 10 μm.

**Supplementary Video 3.** F-actin is enriched within the invasive protrusions at the fusogenic synapse of C2C12 cells Time-lapse imaging of fusion between an F-tractin-mCherry/GFP co-expressing C2C12 cell and a F-tractin-mCherry expressing C2C12 cell at day two in DM on a micropattern. Note that F-actin was enriched in the invasive protrusion (arrowhead) at the fusion site and dissolved immediately after cell fusion, indicated by GFP transfer (arrow) from the invading cell to the receiving cell. The time interval is 1.5 minutes. Single focal plane is shown. Scale bar: 10 μm.

**Supplementary Video 4.** F-actin propelled invasive protrusions mediate the fusion of satellite cells in DM Time-lapse imaging of a fusion event between two mouse satellite cells expressing LifeAct-mNG at day two in DM. F-actin was enriched within the finger-like invasive protrusions at the fusogenic synapse (arrows) and dissolved immediately after cell fusion, indicated by the transfer of LifeAct-mNG from the invading cell to the receiving cell. The time interval is two minutes. Single focal plane is shown. Scale bar: 5 μm.

**Supplementary Video 5.** F-actin and Arp2 are enriched at fusogenic synapse of satellite cells during Erk1/2 inhibition-induced fusion Time-lapse imaging of a fusion event between two mouse satellite cells co-expressing LifeAct-mScar and Arp2-mNG at day two in growth medium (GM) containing 1μM Erk1/2 inhibitor (SCH772984). Note the F-actin and Arp2 enrichment at invasive front (arrowheads) of the invading cell prior to cell-cell fusion and its disappearance after cell-cell fusion. The time interval is two minutes. Max z-projection of 8-10 focal planes from the ventral plasma membrane (z-step size: 500 nm) is shown. Scale bar: 10 μm.

**Supplementary Video 6.** F-actin propelled invasive membrane protrusions mediate the fusion of MyoD^OE^ C2C12 cells. Time-lapse imaging of a fusion event between two C2C12 cells co-expressing LifeAct-mScar and FS-EGFP in GM at day two post MyoD overexpression. Note that F-actin was enriched within the finger-like invasive protrusions at the fusogenic synapse (arrows) and dissolved immediately after cell fusion, indicated by the transfer of FS-GFP from the invading cell to the receiving cell. The time interval is two minutes. Single focal plane is shown. Scale bar: 3 μm.

**Supplementary Video 7.** Migration of NPFs- and Arp2/3 complex-depleted C2C12 cells Time-lapse imaging of sgNT, N-WASP^-/-^, WIP2^-/-^, WAVE2^-/-^, shNT, and Arp2 KD (shArp2) C2C12 cells expressing cytosolic mScar. Note that the mutant cells migrate normally compared to the sgNT or shNT cells. The time interval is five minutes. Max z-projection of 8-10 focal planes from the ventral plasma membrane (z-step size: 3 μm) is shown. Scale bar: 200 μm.

**Supplementary Video 8.** N-WASP, WIP2 and Arp2, but not WAVE2, are enriched at the fusogenic synapse of C2C12 cells Time-lapse imaging of fusion events between C2C12 cells co-expressing LifeAct-mScar and mNG-N-WASP (mNG-WIP2, mNG-WAVE2 or Arp2-mNG,) in GM at day two post MyoD overexpression. Note that N-WASP, WIP2 and Arp2, but not WAVE2, enriched with F-actin prior to fusion, and dissolved after fusion was completed. The sites of LifeAct-mScar enrichment are indicated by arrowheads. The time interval is two minutes. Max z-projection of 8-10 focal planes from the ventral plasma membrane (z-step size: 500 nm) is shown. Scale bar: 5 μm.

**Supplementary Video 9**. **N-WASP is enriched in the shafts of straight protrusions** Time-lapse imaging of protrusion formation in C2C12 cells co-expressing LifeAct-mScar and mNG-N-WASP in GM at day two post MyoD overexpression. Note that N-WASP was enriched with F-actin along the shafts of the protrusions. The enrichment of LifeAct-mScar and mNG-N-WASP on randomly picked protrusions are indicated by green and red arrowheads, respectively. The time interval is one minute. Single focal plane is shown. Scale bar: 5 μm.

**Supplementary Video 10**. **WIP2 is enriched in the shafts of straight protrusions** Time-lapse imaging of protrusion formation in C2C12 cells co-expressing LifeAct-mScar and mNG-WIP2 in GM at day two post MyoD overexpression. Note that, similar to N-WASP, WIP2 was enriched with F-actin along the shafts of the protrusions. The enrichment of LifeAct-mScar and mNG-WIP2 on the protrusions are indicated by green and red arrowheads, respectively. The time interval is one minute. Single focal plane is shown. Scale bar: 5 μm.

**Supplementary Video 11.** N-WASP/WIP2 and WAVE2 are required to generate invasive protrusions Time-lapse imaging of control, N-WASP^-/-^, WIP2^-/-^, WAVE2^-/-^, and Arp2 KD C2C12 cells expressing LifeAct-mNG in GM at day two post MyoD overexpression. Note the straight vs. bendy finger-like protrusions projected from the leading edge of the lamellipodia in the control vs. N-WASP^-/-^ cells; the absence of lamellipodia in the WIP2^-/-^, WAVE2^-/-^, and Apr2 KD cells; the bendy protrusions in the WIP2^-/-^ cell; the decreased number of protrusions in the WAVE2^-/-^ cell; and no protrusion formation in the Arp2 KD cells. The time interval is two minutes. Max z-projections of 8-10 focal planes from the ventral plasma membrane (z-step size: 500 nm) are shown. Scale bar: 10 μm.

**Supplementary Video 12.** WIP2 bundles Arp2/3 complex-mediated branched actin filaments Time-lapse TIRF imaging of actin bundle formation mediated by mNG-WIP2-N and Shi (*Drosophila* homolog of mammalian dynamin). The TIRF assay was performed in NEM-myosin II-coated flow chambers. The time interval is ten seconds.

**Supplementary Video 13.** Visualizing WIP2 bundling actin filaments at the single filament level Time-lapse TIRF imaging of mNG-WIP2-N-mediated bundling of actin filaments at the single-filament level. The TIRF assay was performed with NEM-myosin II-coated flow chambers. mNG-WIP2-N was added into the reaction mix of pre-assembled actin filaments. Left panel shows the bundling of two connected F-actin filaments. Middle and right panels show the bundling of two unconnected F-actin filaments. In each panel, the two pre-bundled actin filaments are indicated by green or red arrowheads, and the resulting bundle is indicated by yellow arrowheads. The time interval is 5 seconds. Scale bar: 2 μm.

**Supplementary Video 14.** The Arp2/3 complex initiates new branched actin polymerization for protrusion initiation and growth Time-lapse imaging of protrusion formation in C2C12 cells co-expressing LifeAct-mScar and Arp2-mNG in GM at day two post MyoD overexpression. Note the Arp2-mNG expanded (red arrowhead) out of the sides of a protrusion, leading to the formation of an actin cloud. Once a new bundle started to form within the actin cloud (green arrowhead), Arp2 continued to polymerize new branched actin filaments at the periphery of the bundle (red arrowheads), which contributed to the growth of the protrusion. The time interval is one minute. Single focal plane is shown. Scale bar: 3 μm.

**Supplementary Video 15.** Dyn2 crosslinks new branched actin filaments into the F-actin bundles during protrusion growth Time-lapse imaging of protrusion growth in C2C12 cells co-expressing LifeAct-mNG and mScar-Dyn2 in GM at day two post MyoD overexpression. Dyn2 was first seen enriched with the actin clouds at the periphery of the F-actin bundle, until the new actin filaments integrated into the actin bundle, making the latter thicker. The actin cloud and mScar-Dyn2 enrichment are indicated by green and red arrowheads, respectively. The time interval is one minute. Single focal plane is shown. Scale bar: 3 μm.

**Supplementary Video 16.** The Arp2/3 complex and Dyn2 are co-enriched during protrusion growth Time-lapse imaging of a growing protrusion in C2C12 cells co-expressing Arp2-mNG and mScar-Dyn2 in GM at day two post MyoD overexpression. Note the colocalization of Arp2 and Dyn2 during protrusion growth. The enrichments of Arp2-mNG and mScar-Dyn2 on the protrusion are indicate by green and red arrowheads, respectively. The time interval is one minute. Single focal plane is shown. Scale bar: 3 μm.

**Supplementary Video 17.** Pharmacological inhibition of Dyn2 impairs the formation of invasive protrusions Time-lapse imaging of C2C12 cells expressing LifeAct-mNG at day two post MyoD overexpression in GM. 5 μM dynamin inhibitor MiTMAB or the empty vehicle (water) were added to the medium 24 hours before the live imaging. Note that the MiTMAB addition led to persistent actin clouds along the protrusions (indicated by arrows), whereas the control cells generated tightly organized actin bundles (indicated by arrowheads). The time interval is one minute. Max z-projection of 8-10 focal planes from the ventral plasma membrane (z-step size: 500 nm) is shown. Scale bar: 5 μm.

**Supplementary Video 18.** Localization of Dyn2 during protrusion formation upon dynamin inhibitor MiTMAB treatment Time-lapse imaging of C2C12 cells expressing LifeAct-mNG and mScar-Dyn2 at day two post MyoD overexpression in GM. 5 μM dynamin inhibitor MiTMAB were added to the medium 24 hours before the live imaging. Note that although Dyn2 was highly enriched in the actin clouds (indicated by red arrowhead) and closely associated with actin enrichment (indicated by green arrowheads), it failed to organize the branched actin filaments into well-defined bundles. The time interval is two minutes. Single focal plane is shown. Scale bar: 5 μm.

**Supplementary Video 19.** The spatiotemporal coordination of Dyn2 and WIP2 during the formation of invasive protrusions Time-lapse imaging of C2C12 cells expressing mScar-Dyn2 and mNG-WIP2 at day two post MyoD overexpression in GM. The mScar-Dyn2 and mNG-WIP2 enrichments are indicated by red and green arrowheads, respectively. Note that mScar-Dyn2 was initially present evenly in the actin cloud (2 min), followed by its enrichment at the periphery of the growing protrusion (4 min), and the subsequent convergence of enriched Dyn2 into the middle of the protrusion (6 min). This was when WIP2 became highly enriched in the actin bundle emerging from the center of the cloud (6 min), and remained enriched in the bundle even after Dyn2’s level decreased. The time interval is two minutes. Single focal plane is shown. Scale bar: 5 μm.

## Materials and Methods

### Animal Ethics

All of the animal studies were approved by the UT Southwestern Medical Center Animal Care and Use Committee according to NIH guidelines.

### Mouse Lines

C57BL/6J (stock: 000664) and Pax3^Cre^ (stock: 005549) mice were obtained from The Jackson Laboratory. The MyoD^iCre^, N-WASP^flox/flox^, CYFIP1^flox/flox^ and ArpC2^flox/flox^ mice were generously provided by Drs. David J. Goldhamer, Scott B. Snapper, Christa W. Habela and Rong Li, respectively. All the mice were housed in 12 light:12 dark cycle at temperatures of 18-23°C with 40-60% humidity. The mouse embryos were staged according to Kaufman^77^. Noon on the day when a vaginal plug was observed was taken as embryonic day 0.5 of gestation (E0.5). The control and mutant littermates were used in each cohort of experiment. Both male and female embryos were used.

### Satellite Cell Isolation and Cell Culture

Satellite cells were isolated from limb skeletal muscles of wild-type C57BL/6J male mice at the age of eight weeks. Briefly, muscles were minced and digested in 800U/ml type II collagenase (Worthington; LS004196) in F-10 Ham’s medium (ThermoFisher Scientific; 11550043) containing 10% horse serum at 37 °C for 90 minutes with rocking to dissociate muscle fibers and dissolve connective tissues. The dissociated myofiber fragments were collected by centrifuge and digested in 0.5U/ml dispase (Gibco; 17105041) in F-10 Ham’s medium at 37°C for 30 minutes with rocking. Digestion was stopped with F-10 Ham’s medium containing 20% FBS. Cells were then filtered from debris, centrifuged and resuspended in satellite cell growth medium (SCGM: F-10 Ham’s medium supplemented with 20% FBS, 4 ng/ ml FGF2 (PeproTech; 100-18B), 1% penicillin–streptomycin and 10mM HEPEs). The cell suspension from each animal was pre-plated twice into the regular 150 mm tissue culture-treated dishes for 30 minutes at 37°C to eliminate fibroblasts. The supernatant containing mostly myoblasts was then transferred into collagen type I-coated dishes and maintained in SCGM.

C2C12 cells (CRL-1772) and U2OS cells (HTB-96) were purchased from ATCC. Tamoxifen-inducible dynamin 1/2/3 triple KO (tKO) mouse embryonic fibroblasts (MEFs) were generously provided by Dr. Pietro De Camilli^62^. All the cells were maintained in growth medium (GM; DMEM supplemented with 10% FBS, 1% penicillin/streptomycin and 10mM HEPEs). For low serum induced muscle cell differentiation, C2C12 cells were cultured in differentiation medium (DM; DMEM supplemented with 2% horse serum, 1% penicillin-streptomycin and 10mM HEPEs). Deletion of dynamin proteins in dynamin tKO MEFs were achieved by supplement of 2 μM 4-hydroxytamoxifen (Sigma; H6278) in the GM for 72 days. Cells in all the conditions were cultured at 37 °C with 5% CO_2_.

### Pharmacological Treatments of C2C12 cells

To pharmacologically inhibit actin polymerization during myoblast fusion, the 100% confluent C2C2 cells were differentiated in DM for 48 hours, followed by incubation with DMSO (0.05%), cytochalasin D (18nM), Arp2/3 complex inhibitor (50 μM) or formin inhibitor SMIFH2 (10 μM) for 24 hours. Then, the cells were fixed in 4% paraformaldehyde (PFA) and immunostained with anti-MyoG and anti-MHC to assess their differentiation and fusion index.

To observe the effect of MiTMAB on the cortical protrusion formation and the localization of Dyn2, LifeAct-mNG or LifeAct-mNG/mScar-Dyn2 expressing C2C12 cells were cultured on fibronectin coated glass coverslips at 30% confluency. After 24 hours, the cells were infected by retrovirus containing MyoD. 24 hours post MyoD overexpression, the cells were cultured in GM containing with MiTMAB (5 μM) or the empty vehicle (water). After another 24 hours, the treated cells were subjected to live imaging.

### Immunocytochemistry

For general immunostaining, cells were fixed in 4% PFA for 15 minutes at room temperature (RT). To immunostain α-actin, β-actin and γ-actin, the cells were fixed in acetone/methanol mixture (volume ratio of 1:1) at −20°C for ten minutes. After extensive washes with PBS, the cells were incubated with blocking buffer (PBS containing 2% BSA and 0.1% TritonX-100) for 20 minutes at RT. Then the cells were incubated with primary antibodies diluted in the blocking buffer at 4°C overnight. After extensive washes with PBS, the cells were further incubated with the appropriate Alexa Fluor-conjugated secondary antibodies diluted in the blocking buffer for one hour at RT. The following primary antibodies were used: rabbit anti-WAVE2 (1:200; Cell Signaling Technologies; 3659), mouse anti-MyoG (1:30; DSHB; F5D), mouse anti-MHC (1:100; DSHB; MF20), mouse anti-Dyn2 (Santa Cruz; sc-166669) and mouse anti-VASP (1:100, Santa Cruz; sc-46668), rabbit anti-Mena (Novus; NBP1-87914), rabbit anti-EVL (Proteintech; 13484-1-AP), mouse anti-α-actin (Sigma; A2172), mouse anti-β-actin (Proteintech; 66009-1-Ig) and mouse anti-γ-actin (Santa Cruz; sc-65635). The Alexa Fluor 488-, 568- and 647-conjugated (Invitrogen) secondary antibodies were used at 1:200. For F-actin labelling, Alexa Fluor 405-, 488-, 568- or 647-conjugated phalloidin (Invitrogen) was used at 1:400 for overnight at 4°C. The cells were then washed with PBS and imaged using a Leica TCS SP8 inverted microscope with Leica Las X software (3.5.5.19976). Images are shown as either single focal planes or maximum z-projection of focal planes as indicated in the figure legends. To compare the protein intensity for different conditions in each experiment, the brightness and contrast levels were set the same for each channel of the comparisons.

### Immunohistochemistry

Mouse embryos were fixed in 4% PFA at 4°C for three days. Whole mount forelimb staining was performed using alkaline phosphatase (AP) conjugated myosin antibody as described in a previous study^78^. Briefly, the skinned forelimbs were post-fixed in 100% methanol prior to the staining. After extensive washes with PBST (PBS containing 0.1% Tween-20), the limbs were incubated in PBST for one hour at 70°C to inactivate endogenous AP, followed by bleaching with 6% Hydrogen Peroxide in PBST for one hour at RT. The bleached samples were then washed in PBST and incubated in the blocking solution (0.1% Triton; 1% BSA; 0.15% glycine in PBS) at RT with rocking. The limbs were subsequently incubated with anti-myosin-AP (My32) (1:800, Sigma; A4335) in blocking solution overnight at 4°C with rocking, followed by extensive washes with PBST. Next, the limbs were washed in NTMT (100mM NaCl, 100mM Tris-HCl (pH 9.5), 50mM MgCl_2_, 0.1% Tween-20) for 15 minutes and developed with NBT/BCIP substrate solution (ThermoFisher Scientific; 34042) at RT.

For immunofluoresence staining of muscle tissue sections, the PFA fixed embryos were dehydrated in 30% sucrose at 4°C overnight. The specimens were embedded in Tissue-Plus O.C.T. Compound (Fisher Scientific; 23-730-571) and 12-μm cryosections were collected onto Superfrost Plus Microscope Slides (Fisher Scientific;12-550-15). Then, the cryosections were incubated with blocking buffer for 20 minutes at RT, followed by overnight incubation with mouse anti-MyoG (1:30; DSHB; F5D) and/or mouse anti-MHC (1:100; DSHB; MF20) at 4°C overnight. After extensive washing with PBS, the sections were incubated with Alexa Fluor-conjugated antibodies for one hour at RT. Subsequently, the sections were washed with PBS and subjected to imaging using a Leica TCS SP8 inverted microscope with Leica Las X software (3.5.5.19976). To compare the protein intensity for different conditions in each experiment, the brightness and contrast levels were set the same for each channel of the comparisons.

### Quantification of Fusion Index

To quantify the cell fusion index, the cell culture was stained with anti-MHC (or phalloidin) and DAPI, and was imaged by epifluorescence microcopy. The fusion index was calculated as the ratio of the nuclei number in myotubes with ≥ 3 nuclei versus the total number of nuclei.

### Quantification of Protrusion Bendiness

The bendiness of the protrusions were quantified as Δtortuosity index, which is the tortuosity index of each protrusion subtracted by the mean tortuosity index of protrusions in Ctrl cells.

### CRISPR sgRNA Knockout in Myoblasts

The sgRNAs targeting the mouse *N-WASP*, *WIP2* and *WAVE2* open reading frames were designed using the online software CRISPOR v5.2. The sgRNAs were individually cloned into lenti-CRISPR v2 vector (Addgene; 52961). The sgRNA sequences are as follows:

N-WASP gRNA, 5’ CCGCGGAGGGTCACCAACGT 3’;

WAVE2 gRNA, 5’ TCCAGCTCGCTTGTATCGCT 3’;

WIP2 gRNA, 5’ AGCACTCCGATCATTAACGT 3’.

The lenti-CRISPR v2 constructs containing specific sgRNAs were transfected into Lenti-X 293T cells (Takara; 632180) with psPAX2 and VSV-G plasmids using the FuGENE HD transfection reagent (Promega; E2311) to produce lentivirus. Two days after transfection, the lentivirus supernatants were filtered and mixed with polybrene (7 μg/ml) to infect the C2C12 cells. Seven days post infection, single cell clones were isolated and expanded. Western blot was used to detect the expression level of target proteins to select the knockout cell clones.

### Retroviral Vector Preparations and Expression

The wild-type or mutant open reading frames of mouse *MyoD*, *MyoG*, *N-WASP*, *WAVE2*, *WIP2* and *Arp2* were amplified using cDNAs generated from C2C12 cells at day two in DM. EGFP, LifeAct, mNeongreen (mNG), mScarleti, Dyn2, and F-tractin-mCherry were amplified from farnesylation signal (FS)-EGFP (Addgene; 21836), LifeAct-mNeongreen (Addgene; 98877), LCK-mScarleti (mScar) (Addgene; 98821), Dyn2-pmCherryN1 (Addgene; 27689) and C1-F-tractin-mCherry (Addgene; 155218), respectively. The wild-type or PRD domain mutated shibire (Shi^4RD^) were descripted previously^17^. For rescue experiments, sgRNA or shRNA-insensitive DNA cassettes for the target genes were generated and used. All the constructs were assembled into the retroviral vector pMXs-Puro (Cell Biolabs; RTV-012) using the NEBuilder HiFi DNA Assembly Cloning Kit (NEB; E2621L). The Sequences of oligonucleotides used for cloning is listed in **Supplementary Table 1**. To package the retrovirus, two micrograms of retroviral plasmid DNA was transfected into platinum-E cells (Cell Biolabs; RV-101) using the FuGENE HD transfection reagent. Two days after transfection, the virus-containing medium was filtered and used to infect cells.

To generate the cell lines expressing cytosolic EGFP, cytosolic mScar, LifeAct-mScar, LifeAct-mNG, F-tractin-mCherry, 3×flag-N-WASP, 3×flag-WIP2, mNG-tagged mutant WIP2 proteins, Shi, Shi^4RD^ or Dyn2-3×HA, retrovirus containing the corresponding construct was filtered, mixed with polybrene (7 μg/ml) and used to infect the cells. One day after infection, cells were washed with PBS and cultured in GM. The EGFP/ F-tractin-mCherry coexpressing cell lines were generated by infection of F-tractin-mCherry expressing C2C12 cells with retrovirus containing EGFP. The Arp2-mNeongreen/LifeAct-mScar-, mNG-N-WASP/LifeAct-mSca-, mNG-WIP2/LifeAct-mScar- and mNG-WAVE2/LifeAct-mScar and mScar-Dyn2/LifeAct-mNG-coexpressing cells were generated by infection of LifeAct-mScar- or LifeAct-mNG-expressing C2C12 cells with retrovirus containing the corresponding construct.

### MyoD- or MyoG-induced Invasive Protrusion Formation and Cell Fusion in Growth Medium

To observe the invasive protrusion formation in the MyoD^OE^ C2C12 cells or MyoD^OE^ MEFs, cells were seeded on fibronectin (Sigma; F1141) coated glass coverslips at 30% confluency in GM. After one day, retrovirus containing MyoD or MyoG was mixed with polybrene (7 μg/ml) and added to GM to infect the cells for a day. Subsequently, the infected cells were incubated in fresh GM for another day (C2C12 cells) or another two days (MEFs) and subjected to live or fixed cell imaging.

To analyze fusion index of the MyoD^OE^ or MyoG^OE^ cells, cells were seeded in tissue culture-treated 6-well plates at 60% confluency in GM. After one day, the cells were infected by retrovirus containing MyoD or MyoG for a day, followed by a fresh GM change. After two (C2C12 cells) or four days (MEFs), the cells were harvested for immunostaining and/or western blot.

### shRNA-based Gene Silence

ShRNAs against *N-WASP*, *WIP2*, *WAVE2* and *Arp2* were individually cloned to lentiviral RNAi construct LENC (Addgene; 111163) or Tet-On lentiviral RNAi construct RT3REN (Addgene; 111166)^78^. The shRNAs against Dyn2 were purchased from Sigma (Mission shRNA).

The shRNA sequences used in this study are as follows:

Sh1N-WASP: TAGTCACACATTTCTTGCCGAG

Sh2N-WASP: TTTTGCTTCTTCTTCATTGGCA Sh1WIP2: TAGTGAACAATTAAAAGTCCGG Sh2WIP2: TTAGTGTGTATGCACTGCTCAT

Sh1WAVE2: TTATCTTTCTTTTCTTTCCTAT Sh2WAVE2: TAACATTATTTGAAGTAGCTAA ShArp2: TTTCTTGGTACTCTTGTCTGGT

Sh1Dyn2 (Sigma; TRCN0000089960): GCCCTTGAGAAGAGGCTATAT Sh2Dyn2 (Sigma; TRCN0000349997): GAGCTCCTTTGGCCATATTAA

The constructs were co-transfected with psPAX2 and VSV-G plasmids into Lenti-X 293T cells using the FuGENE HD transfection reagent to produce lentivirus. Two days after transfection, the medium containing lentiviruses were harvested, mixed with polybrene (7 μg/ml) and used to infect the C2C12 cells. Two days later, puromycin (2 μg/ml) was added to GM for three days to select the infected cells. The surviving cells were subsequently expanded in GM and seeded at 60% confluence in tissue culture-treated 6-well plates. For straight knockdown experiments, when cells grew to 100% confluence, the GM was switched to DM. The cells were then allowed to differentiate for five days during which their differentiation and fusion were assessed. For doxycycline-induced shRNA expression, doxycycline (1 μg/ml) or empty vehicle was added to GM 24 hours after the cells were seeded. 48 hours post doxycycline addition, the GM was switched to DM with doxycycline (1 μg/ml) or empty vehicle. The cells were then allowed to differentiate for five days during which their differentiation and fusion were assessed.

### Western Blot

For western blot, cells were lysed by ice-cold RIPA buffer (150mM NaCl, 1% NP40, 0.1% SDS and 50mM Tris, PH7.4) containing protease and phosphatase inhibitor (Cell Signaling Technologies; 5872) for 20 minutes. The supernatants were collected by centrifugation at 140,000 x g for 15 minutes. Protein concentrations were determined using the Bradford Protein Assay Kit (Bio-Rad; 5000201). 10-30μg total protein was loaded and separated by 10% SDS-PAGE gel and transferred to PVDF membranes (Millipore; GVHP29325). Then, the membranes were blocked for one hour at RT in PBS containing 5% nonfat dry milk and 0.1% Tween-20 (PBSBT) and subsequently were incubated with primary antibodies diluted 1:1000 in PBSBT overnight at 4°C. The membranes were then washed with PBST and incubated with appropriate HRP-conjugated secondary antibodies diluted in PBSBT for one hour at RT. After extensive washes with PBST, the membranes were developed with the ECL western blotting substrate (ThermoFisher Scientific; 32209). The following primary antibodies were used: mouse anti-WASP (1:1000; Santa Cruz; 4860), rabbit anti-N-WASP (1:1000; Cell Signaling Technologies; 4848), mouse anti-WIP1 (1:1000, Santa Cruz; sc-271113), rabbit anti-WIP2 (1:1000; ThermoFisher Scientific; PA5-55086), mouse anti-WAVE1 (1:1000; Santa Cruz; sc-271507), rabbit anti-WAVE2 (1:1000; Cell Signaling Technologies; 3659), rabbit anti-WAVE3 (1:1000; Cell Signaling Technologies; 2806), mouse anti-Arp2 (1:1000; Santa Cruz; sc-166103), mouse anti-Arp3 (1:1000; Santa Cruz; sc-48344), mouse anti-MyoG (1:30; DSHB; F5D), mouse anti-MHC (1:200; DSHB; MF20), rabbit anti-ABI1 (1:1000; Cell Signaling Technologies; 39444), rabbit anti-CYFIP1(1:1000; Cell Signaling Technologies; 81221), mouse anti-mNeongreen (1:2000; Proteintech; 32F6), mouse anti-actin (Sigma: A4700), sheep anti-ESGP (1:1000; R&D Systems; AF4580), mouse anti-α-actin (1:50000; Sigma; A2172), mouse anti-β-actin (1:50000; Proteintech; 66009-1-Ig), mouse anti-γ-actin (1:50000; Santa Cruz; sc-65635), mouse anti-Dyn2 (1:1000; Santa Cruz; sc-166669) and rabbit anti-β-Tubulin (1:1000; Cell Signaling Technologies; 2146).

### Protein Purification

Three partial sequences of mouse WIP2, aa1-176 (WIP2-N), aa35-53(WH2-1) and aa80-105 (WH2-2), were fused with mNeongreen at their N-terminus with a flexible peptide linker and cloned into pET28a vector (Millipore; 69864). Protein expression in *Escherichia coli* BL21(DE3) cells was induced by 200 μM IPTG (OD600=0.8) for 18 hours at 16°C. Bacteria were collected and lysed by sonication in lysis buffer (50 mM Tris-HCl pH 8.0, 500 mM NaCl, 5 mM imidazole, 5 mM β-mercaptoethanol) supplemented with protease inhibitors, followed by centrifugation at 20,000 rpm for 45 minutes at 4°C. The proteins were then purified with Ni-NTA agarose beads (QIAGEN; 30210) according to the manufacturer’s instructions. Proteins eluted by 100 mM imidazole were collected and then desalted with disposable PD 10 desalting columns (Sigma, GE17-0851-01) against stocking buffer (20 mM HEPES (pH 7.3), 150 mM KCl, 1 mM EGTA and 1 mM dithiothreitol). The purified proteins were subjected to SDS-PAGE gels and coomassie blue staining to determine the purity of the proteins. Bradford assays were performed to determine the protein concentrations. The purified proteins were then flash frozen in liquid nitrogen and stored at −80 °C until use.

### F-actin Co-sedimentation Assay

F-actin co-sedimentation assays were performed to determine the F-actin binding and bundling activities of mouse WIP2. F-actin pre-assembled from 5 μM G-actin was incubated alone or with the indicated concentrations of proteins in 5 mM HEPES (pH 7.3), 50 mM KCl, 2 mM MgCl_2_, 0.5 mM EGTA, 3 mM imidazole, 60 nM ATP, 60 nM CaCl_2_, 0.4 mM DTT and 0.003% NaN_3_ for 30 minutes at RT. The samples were then spun for 30 minutes at 4 °C at 100,000 x g (high-speed co-sedimentation) or 13,600 x g (low-speed co-sedimentation). After centrifugation the supernatants and pellets were resolved by 10% SDS-PAGE and the gels were stained with coomassie blue or subjected to western blot.

### Pharmacological Treatments of Dynamin-actin Interaction *in vitro*

To determine the effect of dynamin inhibitor, MiTMAB, on dynamin-actin interaction *in vitro*, F-actin pre-assembled from 1 μM G-actin was incubated alone or with 1μM His-Shibirie in 5 mM HEPES (pH 7.3), 50 mM KCl, 2 mM MgCl_2_, 0.5 mM EGTA, 3 mM imidazole, 60 nM ATP, 60 nM CaCl_2_, 0.4 mM DTT and 0.003% NaN_3_ for 30 minutes at RT. Subsequently, 2mM GTP with or without 40 μM MiTMAB was added to the appropriate reaction mixtures and incubated for five minutes at RT. Then, the samples were mixed with Alexa Fluor 488-conjugated phalloidin (Invitrogen), dropped on the glass coverslips, and subjected to imaging using a Leica TCS SP8 inverted microscope with Leica Las X (3.5.5.19976).

### Time-lapse Imaging and Analysis

Time-lapse imaging was performed with a Nikon A1R confocal microscope with NIS-Elements AR (5.10.01 64-bit) at 37 °C and 5% CO_2_ using a Nikon Biostation CT. The time interval of imaging for each video is described in the supplementary video legends. After time-lapse imaging, ImageJ (NIH, 64-bit 1.52f) was used to assemble the images into time-lapse videos.

To image C2C12 fusion in DM, cells were seeded on micropatterns (with a diameter of 250 μm) and imaged using a 40× (0.4 NA) objective at 48 hours after switching from GM to DM. The micropatterns were generated as described previously^79^. To quantify the fusion of SiR-Actin^+^ and SiR-Actin^-^ cells, cells on micropattern at 48 hours in DM were incubated with SiR-Actin for 30 minutes to label F-actin, and subjected to live imaging by confocal microscopy for 16 hours.

To image MyoD^OE^-induced fusion and protrusion formation of C2C12 cells cultured in GM, cells were seeded on the fibronectin-coated cover glass and imaged using a 40× (0.4 NA) objective at 48 hours post MyoD overexpression.

To image Erk1/2 inhibition-induced satellite cell fusion, low passage (≤ 3 passages) satellite cells were plated at 50% confluence on collagen type I-coated 6-well plates in SCGM for 24 hours. The following day, the cells were infected by a mixture of retroviruses containing LifeAct-mScar and Arp2-mNG (1:1 in volume) in SCGM. After 48 hours, the cells were trypsinized and plated at 40% confluence on fibronectin-coated cover glass in regular GM containing 1μM ERK1/2 inhibitor (Cayman Chemicals; SCH772984). Subsequently, the cells were subjected to live imaging at 48 hours post SCH772984 treatment 40× (0.4 NA) objective.

To image fusion of satellite cells in DM, LifeAct-mNG expressing low passage satellite cells were generated by infection of retrovirus containing LifeAct-mNG in SCGM. After 48 hours, the cells were trypsinized and plated at 40% confluence on fibronectin-coated cover glass in DM. The cells were subjected to live imaging at 48 hours post DM treatment 40× (0.4 NA) objective.

To examine the C2C12 cell migration, cytosolic mScar expressing C2C12 cells were seeded in tissue culture-treated 6-well plates and imaged in GM using a 10× (0.5 NA) objective.

### Total Internal Reflection Fluorescence (TIRF) Microscopy

Flow chambers were assembled and coated with 10 nM NEM-myosin II in high salt TBS buffer (50 mM Tris-HCl (pH 7.5) and 600 mM NaCl) for one minute, washed with high salt BSA (1% BSA, 50 mM Tris-HCl (pH 7.5) and 600 mM NaCl) and low salt BSA (1% BSA, 50 mM Tris-HCl (pH 7.5) and 150 mM NaCl) sequentially, then washed with the TIRF buffer (50 mM KCl, 1 mM MgCl_2_, 1 mM EGTA, 10 mM imidazole, 100 mM DTT, 0.2 mM ATP, 15 mM glucose, 20 μg/ml catalase, 100 μg/ml glucose oxidase, 0.2% BSA and 0.5% methylcellulose (pH 7.0)). To visualize WIP2 and dynamin-mediated bundling of the Arp2/3 complex-nucleated branched actin network, 50 nM Arp2/3 complex (Cytoskeleton; RP01P), 1 μM VCA domain of WASP (Cytoskeleton; VCG03-A), 2 μM G-actin (70% unlabeled and 30% rhodamine-G-actin; cytoskeleton) were mixed in TIRF buffer, loaded into the flow chambers and subjected to Ring-TIRF imaging immediately using a GE OMX-SR super-resolution microscope equipped with a 60X/1.49 UPlanApo oil objective lens, with AcquireSR (4.5. 10204-1). After 20 minutes, 4 μM mNG-WIP2-N or His-Shibire was loaded into the flow chambers while the actin filaments were continuously imaged by Ring-TIRF.

### Super-Resolution Microscopy

To visualize the distribution of WIP2 along actin bundles, 1 μM mNG-WIP-N was incubated with 1 μM F-actin and 1 μM Alexa Fluor 568-phalloidin in 5 mM HEPES (pH 7.3), 50 mM KCl, 2 mM MgCl_2_, 0.5 mM EGTA, 3 mM imidazole, 60 nM ATP, 60 nM CaCl_2_, 0.4 mM DTT and 0.003% NaN_3_ for 30 minutes at RT. Subsequently, the samples were diluted 20-fold in the fluorescence buffer (0.5% methycellulose, 50 mM KCl, 1 mM MgCl_2_, 1 mM EGTA, 10 mM imidazole, 100 mM DTT, 0.02 mg/ml catalase, 0.1 mg/ml glucose oxidase, 15 mM glucose, 0.1 mM CaCl_2_, 0.1 mM ATP and 0.005% NaN_3_). Then, 4 μl of diluted samples were mounted onto glass slide, covered with a poly-L-lysine-coated coverslip, and subjected to structured illumination microscopy (SIM) using a GE OMX-SR super-resolution microscope equipped with a 60X/1.42 UPlanApo oil objective lens with AcquireSR (4.5.10204-1).

To visualize the distribution of N-WASP/WIP2 and WAVE2 on the invasive actin structures, 3×flag-N-WASP or 3×flag-WIP2 C2C12 cells were immunostained with mouse anti-Flag (1:200; Sigma; F1804), rabbit anti-WAVE2 (1:200; Cell Signaling Technologies; 3659) and Alexa Fluor 647 phalloidin (1:500; ThermoFisher Scientific; A30107) overnight at 4 °C. Subsequently, the cells were substantially washed with PBS and incubated with the Alexa Fluor 488- and 568-conjugated (1:200; Invitrogen) for one hour at RT. After extensive washes with PBS, the samples were subjected to SIM using a GE OMX-SR super-resolution microscope equipped with a 60X/1.42 UPlanApo oil objective lens with AcquireSR (4.5.10204-1).

Stimulated emission depletion microscopy (STED) was used to examine the diameter of invasive protrusions and filopodia. At day two post MyoD overexpression, the wild-type C2C12 cells were immunostained with mouse anti-VASP (1:100, Santa Cruz; sc-46668) and Star Red-conjugated Phalloidin (1:200; Abberior; STRED-0100) overnight at 4 °C. Next, the cells were extensively washed in PBS and incubated with Star Orange-conjugated goat anti-mouse antibody (1:200; Abberior; STORANGE-1001) for one hour at RT. STED imaging was performed using an Abberior Instruments Expert Line STED microscope (Abberior Instruments) equipped with an SLM-based easy3D module, an Olympus IX83 microscope body and an UPLSAPO 60×/1.4 oil objective, with Imspector (16.3). To compare the phalloidin signal intensity in different type of protrusions, the brightness and contrast levels are set the same for each channel of the comparisons.

### Immunolabeling of Dynamin 2 in Membrane Extracted Cells

To visualize Dyn2 on the cytoskeleton of MyoD^OE^ cells, cell membrane was detergent-extracted for immunofluorescence labeling of Dyn2, or immunogold labeling of Dyn2 followed by platinum replica electron microscopy (PREM).

Dyn2 KD C2C12 cells were seeded at 30% confluency in GM on fibronectin coated cover glass. After 24 hours, retrovirus containing MyoD and Dyn2-3×HA was mixed with polybrene (7 μg/ml) and added to GM to infect the cells. After two days post infection the cells were subjected to membrane extraction as described previously^63^. Briefly, the cells on glass coverslips were rinsed twice in PBS++ and incubated in extraction solution containing 1% Triton X-100, 2% polyethylene glycol (or PEG), 2 µM unlabelled phalloidin (Sigma–Aldrich; P2141) and 2 µM taxol (Sigma–Aldrich; T7402) in PEM buffer (100 mM PIPES−KOH (pH 6.9), 1 mM MgCl_2_ and 1 mM EGTA) for 4 min.

Detergent-extracted cells were fixed with 2% glutaraldehyde (Ted Pella; 18426) prepared in PEM buffer at room temperature for 30 min. The fixed cells were quenched with 2 mg/ml NaBH_4_ in PBS twice for 10 min each and blocked with 1% BSA in PBS for 20 min. The cells were then incubated with rabbit anti-HA (Sigma–Aldrich; H6908) antibody diluted 1:100 in PEM buffer for 1 hour, followed by either immunofluorescence or immunogold labeling.

For immunofluorescence labeling of Dyn2, the samples were washed in blocking buffer containing 1% BSA in PBS and incubated with goat-anti-rabbit Alexa Fluo 405 (ThermoFisher Scientific; A-31556) and Alexa Fluo 568 Phalloidin (ThermoFisher Scientific; A12380) diluted (1:300) in blocking buffer overnight. The cells were then washed with PBS and imaged using a Leica TCS SP8 inverted microscope. Images are shown as single focal planes as indicated in the figure legends. The brightness and contrast levels are set the same for each channel of the images.

To perform immunogold labeling of Dyn2 and PREM, the rabbit anti-HA incubated samples were washed in immunogold buffer containing 0.5 M NaCl, 0.05% Tween 20 and 0.05% sodium azide in 20 mM Tris-HCl (pH 8.0) and incubated with secondary antibodies conjugated with 18-nm colloidal gold (Jackson ImmunoResearch; 111-215-144) diluted (1:10) in immunogold buffer overnight. After washing, the samples were post-fixed with 2% glutaraldehyde. For platinum replica, the cells were post-fixed by sequential treatment with 0.1% tannic acid and 0.2% uranyl acetate in water, dehydrated through a graded ethanol series (10, 20, 40, 60 and 80% for 5 min each, 100% twice for 5 min each, 0.2% uranyl acetate in 100% ethanol for 20 min, and 100% ethanol four times for 5 min each), critical point dried, coated with platinum and carbon and transferred to electron microscopy grids for observation. PREM samples were examined using a JEM-1400Plus transmission electron microscope (JEOL USA) operated at 120 kV. Images were acquired using an AMT BioSprint 16M-ActiveVu CCD camera (AMT) and presented in inverted contrast.

### Electron Microscopy

To observe the invasive protrusions at the contact sites of C2C12 cells in DM, C2C12 cells were first cultured on tissue-culture treated 35mm dishes in GM. Once the cells reached 100% confluence, GM was replaced with DM. After 48 hours, the cells were fixed in a solution containing 2.5% glutaraldehyde, 1% sucrose and 3mM CaCl_2_ in 0.1M sodium cacodylate buffer (pH 7.4) at 4°C overnight. On the following day, samples were rinsed in 0.1M cacodylate buffer containing 3% sucrose and 3mM CaCl_2_ before being post-fixed for 1.5 hours on ice with 1% osmium tetroxide and 0.8% potassium ferricyanide. En bloc staining was performed by treating the samples with 0.25% tannic acid and 4% uranyl acetate. Then, the samples were dehydrated, infiltrated, and embedded in EPON resin. Using a LEICA ultramicrotome (UC6), resin-embedded samples were cut into 70 nm sections and collected on copper slot grids. These sections were viewed with a JEOL 1400 transmission electron microscope after being post-stained with 2% uranyl acetate and Reynold’s stain. To observe the invasive protrusions at the contact sites of MyoD^OE^ cells, C2C12 cells cultured on tissue-culture treated 35mm dishes were infected with retrovirus carrying MyoD in GM. 48 hours post infection, the cells were subjected to fixation, staining and electron microscopy as described above.

To observe the invasive protrusions at the contact sites of myoblasts *in vivo*, hindlimbs of E15.5 mouse embryos were fixed in a solution containing 3% paraformaldehyde, 2% glutaraldehyde, 1% sucrose, 3mM CaCl_2_ in 0.1M sodium cacodylate buffer (pH 7.4) at 4°C overnight. Samples were subsequently washed with 0.1M cacodylate buffer containing 3% sucrose and 3mM CaCl_2_, and post fixed with 1% osmium tetroxide in 0.1M sodium cacodylate buffer for 1.5 hours on ice. The limbs were stained with 2% uranyl acetate, dehydrated and embedded in EPON resin. The embedded samples were then cut into 70 nm thick sections using LEICA ultramicrotome (UC6) and collected on copper slot grids. These sections were post-stained with 2% uranyl acetate and sato’s lead solution and examined using a JEOL 1400 transmission electron microscope.

### Negative-Stain Electron Microscopy

Negative-stain electron microscopy was performed to visualize the distribution of WIP2 and dynamin along F-actin bundles. A total of 5 μM F-actin was incubated with an appropriate concentration of mNeogreen-WIP2 or His-dynamin in 6.5 mM HEPES (pH 7.3), 50 mM KCl, 2 mM MgCl_2_, 0.3 mM EGTA, 20 nM ATP, 20 nM CaCl_2_, 0.3 mM DTT and 0.001% NaN_3_ at RT for 30 minutes. The reaction mixture was loaded onto carbon-coated, glow-discharged 400 mesh copper grids and incubated for five minutes, followed by two quick washes in 1 x actin polymerization buffer (50 mM KCl, 2 mM MgCl_2_, 5 mM guanidine carbonate (pH 7.5), and 1 mM ATP). The grid was stained in two drops of 0.75% uranyl formate for 20 seconds and air dried. Electron micrographs were collected on a JEOL 1400 transmission electron microscope.

### Statistics and Reproducibility

Statistical significance was performed using GraphPad Prism 8 software with two-tailed student’s t-test. Sample size and replicates are indicated in the figure legends. All experiments were repeated in at least three independent biological replicates. Precise P values are in the figures. P values < 0.05 were considered statistically significant. The investigators were not blinded to allocation during the experiments and outcome assessment. No data were excluded from the analyses. For the in vivo studies, age-matched embryos were randomly assigned to experimental and control groups. No statistical methods were used to pre-determine sample sizes, but our sample sizes are similar to those reported in previous publications^17,32,33,80^. Data distribution was assumed to be normal but this was not formally tested.

## Data availability

All the data supporting the findings of this study are available within the article, its supplementary information files and the source data files.

